# Dynamical Systems Models for Plasma Dilution

**DOI:** 10.1101/2022.03.15.484505

**Authors:** Dylan Chambers, Neil Christensen

## Abstract

Recent experiments provide evidence that diluting the blood plasma restores the plasma environment to a more youthful level at least partially restoring the health of organs and tissues throughout the body. We propose that a dynamical-systems model representing the plasma constituents could support the optimization process and help determine the appropriate dilution level, frequency and any simultaneous plasma infusions to achieve the most favorable outcome. We use a combination of a gradient descent, a simulated annealing and a genetic algorithm to find a population of models that fit illustrative data. We analyze this population and present distributions of the model parameters and include a collection of plots of the dilution process for illustrative models. We then consider modifications of the dilution in order to illustrate what predictions might be possible had we more data to disambiguate the model.

---

In a stunning pair of experiments, Mehdipour et. al. [1, 2] showed that simply diluting the blood plasma with a combination of saline and 5% albumin, which they called neutral blood exchange (NBE), was sufficient to rejuvenate all three germ layers, improve cognition and attenuate neuroinflammation in old mice. In fact, they found that a single NBE treatment, replacing 50% of the plasma in old mice, resulted in enhanced muscle repair, reduced liver adiposity and fibrosis and increased hippocampal neurogenesis to atleast the level seen in earlier heterochronic parabiosis experiments [3] and surpassing the level seen in earlier heterochronic blood exchange experiments [4]. Moreover, they showed that a single NBE treatment surpassed ABT 263, a senolytic that does not pass the blood-brain barrier, in its rejuvenative effect on the brain.

Additionally, the authors found that the protein composition of the plasma was reset to a more youthful, more rejuvenative composition and, although in this more youthful plasma some proteins were reset to a lower level, as might be expected from a dilution experiment, perhaps surprisingly, many were elevated as a consequence of the NBE treatment. They noted that this was evidence that some of the age-elevated protein factors suppressed the age-reduced protein factors and that once those suppressive proteins were diluted, the inhibited proteins were allowed to upregulate to a higher, more youthful level.

Noting that an equivalent procedure in humans, therapeutic plasma exchange (TPE), has already been approved for treatment by the US Food and Drug Administration (FDA), they further measured the blood proteomics of aged humans before and after a single TPE treatment and found that the protein composition reset in an analogous way to the old mice. The resetting and rejuvenating effect of TPE was further summarized in [8]. Although TPE appears capable of inducing a resetting of the plasma composition, it is not yet fully clear how much plasma must be removed in order to see this resetting. The NBE experiments outlined above replaced approximately 50% of the plasma with saline and albumin while typical TPE treatments replace the entire plasma volume. To begin to answer this, the authors of [9] analyzed human plasma before and after a short series of standard commercial plasma donations without the addition of albumin. They found that some of the same proteins that were enhanced to youthful levels in [1] were also increased as a result of plasma donation. However, others were not significantly altered. A much larger, more comprehensive double-blind, placebo controlled clinical trial is clearly needed to establish the optimal treatment protocol as well as the level of rejuvenation of tissues in humans that results.

Before continuing, we should also note that some researchers have considered the rejuvenating effect of some factors in young plasma and have performed some experiments where, rather than diluting plasma, they infuse young factors into the elderly [10–14]. Of course, both may have merit and the optimal protocol could very well involve a dilution combined with an infusion.

Beyond the measurement of the rejuvenative phenotype and the plasma proteomics, Mehdipour et. al. [1] also built a dynamical-system toy model that illustrated how a dilution might cause an enhancement to some plasma species in the system. This illustrative model had three species with the following set of differential equations (DEs),

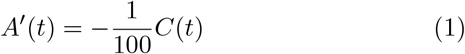

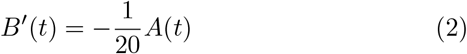

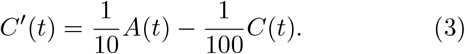

We can see that in their toy model, species *A* positively affected *C* but negatively impacted *B*, while *C* negatively affected *A* and itself (which could be interpreted as its own decay). Schematically, this can be written as,

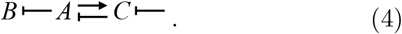

With this model, beginning with the initial values

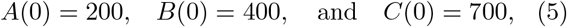

which they took to be diluted from *A*_*BD*_ = 1000, *B*_*BD*_ = 400 and *C*_*BD*_ = 700, where the subscript *BD* stands for before dilution. Using this set of differential equations and initial values, we reproduce their Fig. 6 here, in our Fig. 1. As can be seen, this set of differential equations gives interesting dynamics illustrating the relative values of some species decreasing (*A* and *C*) while another increases (*B*). It also displays oscillations on the way to equilibrium as the species overshoot before the feedback loops and decays remove the oscillatory energy and it reaches equilibrium.

**FIG. 1.**
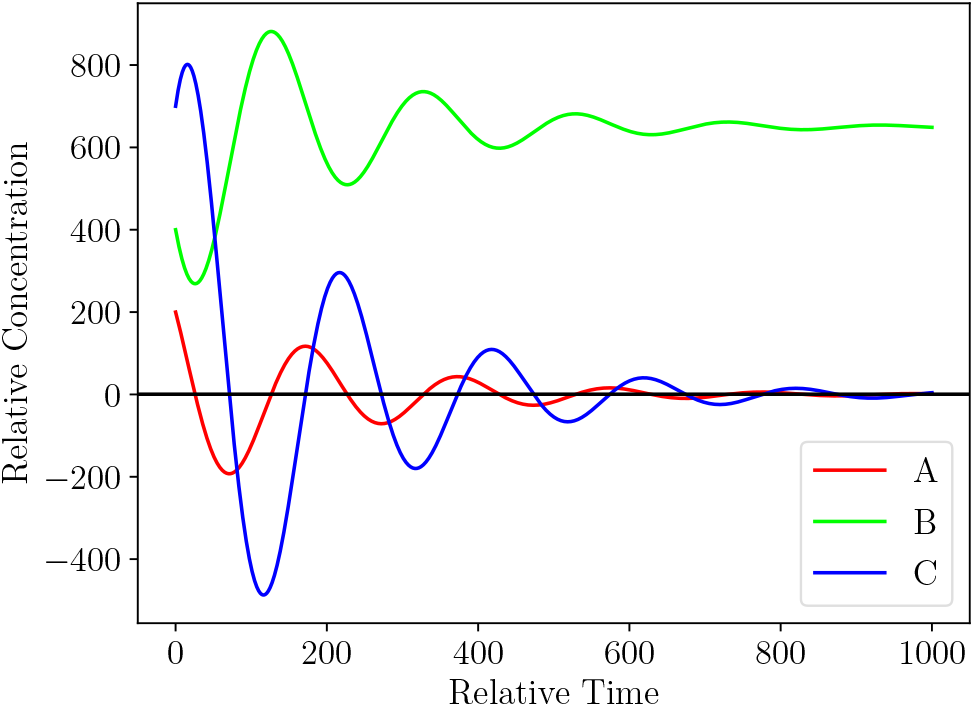
A reproduction of Fig. 6 from [1] using the system of differential equations in Eqs. (1) through (3) and initial values given in Eq. (5).

Although the toy model achieved the illustrative purposes of [1], we would like to improve on their model in a few important ways. We first note that their model only contained one steady state for *A* and *C* and, therefore, the proposed initial state before the dilution with *A*_*BD*_ = 1000, *B*_*BD*_ = 400 and *C*_*BD*_ = 700 does not correspond to a steady state of their model. In fact, a steady state for this system occurs when all the derivatives are zero and the right-hand side of Eqs. (1) through (3) vanish. This only happens when *A* = *C* = 0. On the other hand, *B* can take any steady state value and depends on the initial conditions. It “freezes out” when *A* vanishes. To improve on this, we would like to find models with at least two non-trivial steady states representing the before-dilution and after-dilution states of the system.

Another shortcoming of this toy model is that the species concentrations for *A* and *C* become negative before reaching the vanishing endpoint. This is clearly unphysical and we would like to enforce that the model species never become negative in a more realistic system. In order to attempt an exploration of better models, we designed a system of differential equations as described in App. A and wrote a computer code as described in App. B. We used our code to generate a population of models that fit the following data,

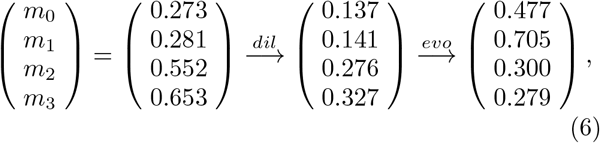

where *dil* stands for dilution and *evo* stands for evolution. These values are meant to be illustrative with *m*_0_ and *m*_1_ increasing to a higher value after dilution and *m*_2_ and *m*_3_ decreasing to a lower value after dilution. We modeled these values on the proteins ROBO4, Angiogenin, VEGF-C and Epiregulin, from Fig. 5E from [1]. However, these values are only used in a demonstrative way, in order to show what might be possible had we a robust dynamical model for plasma dilution. As we will see, we do not yet have enough data to disambiguate the many models that would fit the data.

The remainder of this article is organized in the following way. In Sec. I, we describe the statistical properties of a population of minimal models which only contain the “measured” species in Eq. (6) and no extra species beyond these measured species. In Sec. II, we consider modified dilutions in the models described in Sec. I. That is, we consider other dilution scenarios such as flat dilutions at different values, flat dilutions with a single or two species enhanced, and other possibilities. In Sec. III, we summarize our findings and conclude. We also include several appendices. In App. A, we briefly review dynamical models and describe the structure and parameters of the models used in our analysis. In App. B, we describe our algorithms for generating a population of models that fit the data. In App. C, we present plots illustrating the behavior of the minimal models following the standard dilution (the models and dilutions analyzed in Sec. I). In App. D, we show plots illustrating the behavior of one of the minimal models under different modified dilution scenarios.

## I. MINIMAL MODELS

In this section, we consider minimal models with only the measured species but none of the extra species described in App. A. A selection of the models used for the analysis in this section can be found in App. C. Although the genetic algorithm attempted to add extra species (of the type *u*_*i*_, *h*_*i*_ and *c*_*i*_ – see App. B), none of the models with extra species had low enough model scores to remain in the population. However, it is of course possible for models with extra species to fit the data as well or better than these minimal models. In fact, in a separate calculation, we forced the model to retain at least 4 extra species and found many models that fit the data as well or better than the present minimal models. Nevertheless, we chose to present these minimal models because, firstly, they are minimal and no extra species were required and, secondly, since the models with extra species are more complex and have a much larger space of parameters to search, we were unable to obtain a large enough population of extra-species models with sufficiently low model scores that we could have confidence in the statistical properties of the distributions.

Each new minimal model took a little over 6 cpu-minutes to optimize, on average (some took much longer, some took much shorter). We generated a total of approximately 350 thousand new models which took a total of approximately 1500 cpu-days of computation. We kept the 1000 models with the best model scores (see App. B). In Fig. 2, we show a distribution of the model scores in the final population. The maximum model score in the population depends on how many models we try. Greater run times produces models with lower model scores and lower population maximum score. In the future, with much greater data (especially data with more temporal measurements and data with a variety of dilution values), we would be most interested in only the few very best model scores and ignore all the rest. In the absence of sufficient data to disambiguate the many possible models shown here, we instead focus on traits shared by the many models that fit the data.

**FIG. 2.**
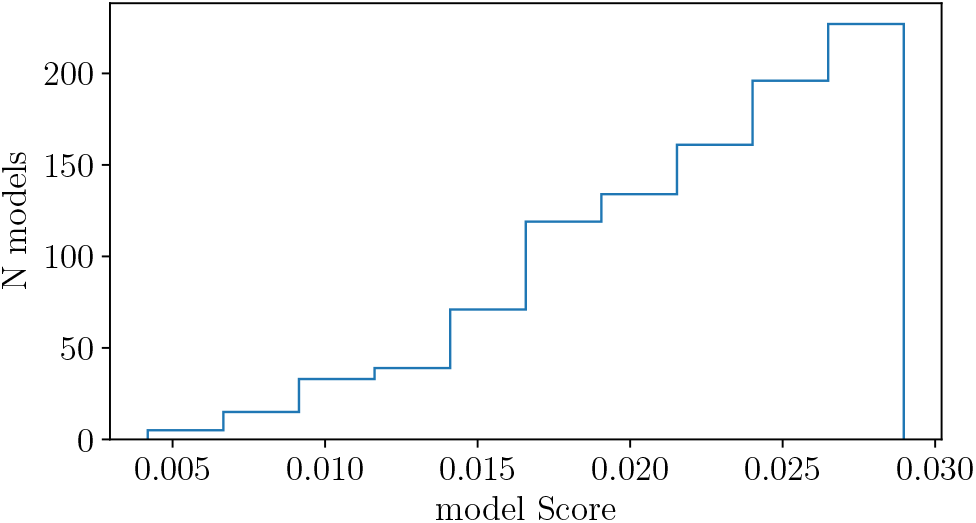
A distribution of model scores from our best 1000 models with only 4 measured species.

As described in App. B, for each species’ equations, when creating a new model from scratch, we randomly chose (with a flat distribution) between 2 and 7 terms [the number of terms on the right of Eq. (A6)], including the final decay term. However, the genetic algorithm sometimes took a pre-existing model from the population and either added a term or removed a term (see App. B), allowing the number of terms to exceed these bounds and change the shape of the distribution. Therefore, the final shape of the distributions is a combination of the flat distribution convoluted with the mating, the mutations and the survival of the fittest algorithms. With this in mind, we present a distribution for the number of terms in the equation for each constituent in the top plot of Fig. 3. We can see that the number of terms per equation is peaked at 2 terms for *m*_0_ and *m*_2_, at 3-4 terms for *m*_1_ and at 4-5 terms for *m*_3_. As expected, there were no models with only 1 term (only the decay term) on the right of any equation, since the steady state for such a species would necessarily be zero. There must be something to balance the decay in order for the species to be nontrivial. After the dilution, *m*_2_ rises a very small amount, which apparently only requires one additional term, *m*_0_ increases a moderate amount, which also only requires one additional term, but *m*_1_ increases twice as much as *m*_0_ and apparently needs at least two or three additional terms. On the other hand, *m*_3_ decreases following dilution, the opposite of the other three and this requires three to four additional terms. Furthermore, as we can see in the middle of Fig. 3, typically only the decay term is negative for *m*_0_ and *m*_2_. Once again, this must be the case, else *m*_0_ and *m*_2_ would be trivial with both terms negative and no non-trivial steady state. On the other hand, one or more of the extra terms is typically negative for *m*_1_ and *m*_3_, presumably due to the extra complexities of their evolutions. For each species’ equation, we have also analyzed how many of their terms are of the form Eq. (A2), with only a single species, or of the form Eq. (A4), with two species and show a plot in the bottom of Fig. 3. Once again, *m*_0_ and *m*_2_ are similar with around 80% being single-species terms, while *m*_1_ has approximately half of its terms with two species and approximately 80% of the terms for *m*_3_ have two species.

**FIG. 3.**
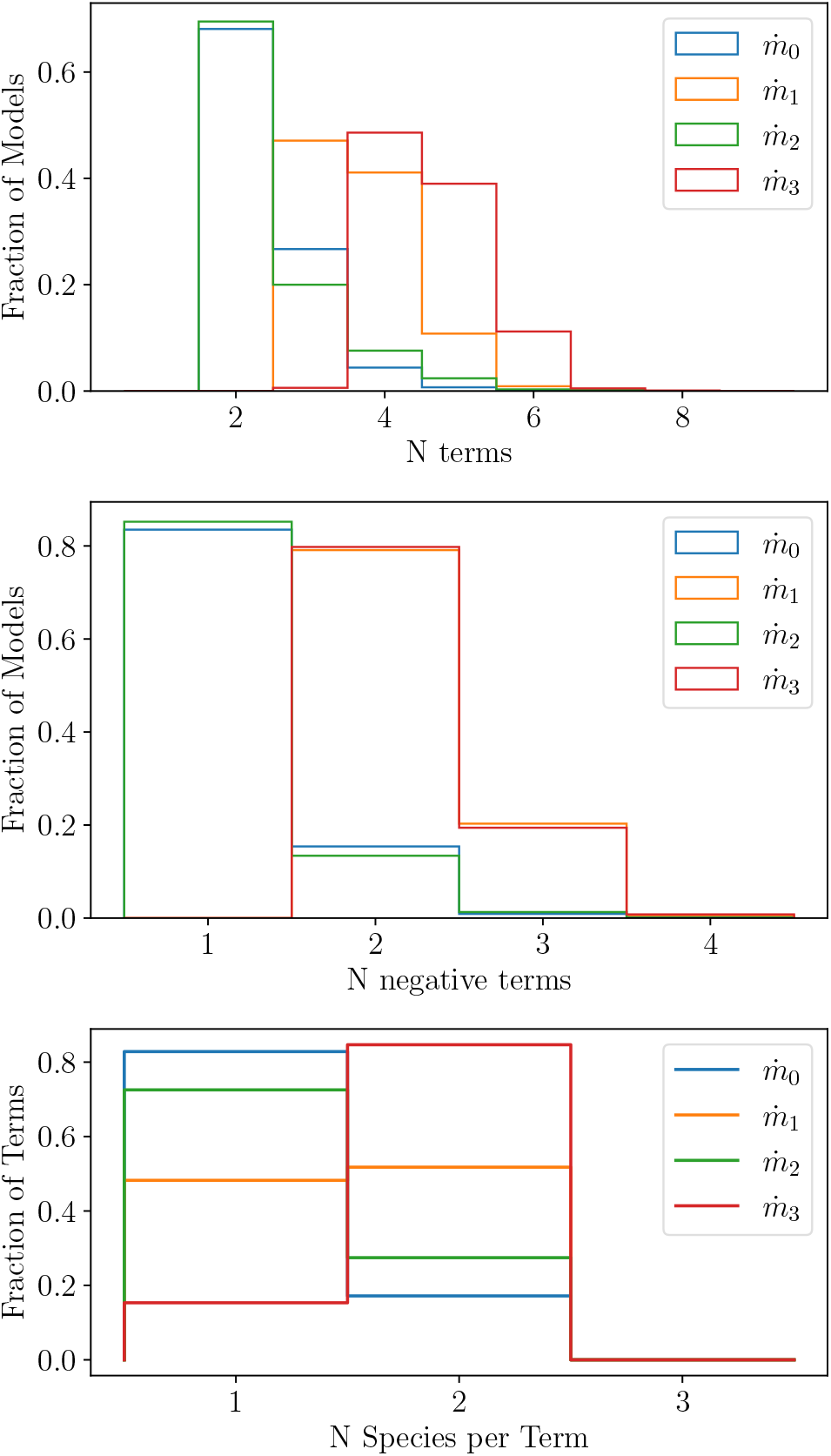
These plots show the fraction of models or terms that have certain properties in the equations for each species. The blue, orange, green and red lines represent the equations for *m*_0_, *m*_1_, *m*_2_ and *m*_3_, respectively. The top plot is for the number of terms per equation. The middle plot is for the number of negative terms per equation. The bottom plot is for the number of species per non-decay term for each equation. In other words, whether a term was of the form Eq. (A2) or Eq. (A4).

If we look at each equation in greater detail, we begin to see further patterns. In Fig. 4, we plot which species is included for single-species terms for each equation. These are terms of the form Eq. (A2). The blue line is for the first equation that gives 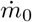. We see that a little over 80% of its single-species terms contain *m*_1_ and a little more than 10% contain itself. The orange line, on the other hand, represents the equation for 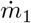 and nearly 70% of its single-species terms contain *m*_0_, while nearly 30% contain itself. It makes sense that these two are tightly coupled since they share the property that both increase following dilution to a steady state that is higher than the previous steady state. As we can see, species *m*_2_ and *m*_3_ are similarly related for single-species terms. We see from the green line that approximately 90% of the single-species terms for 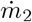 contain *m*_3_ while approximately 80% of the single-species terms for 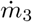 contain *m*_2_, in red.

**FIG. 4.**
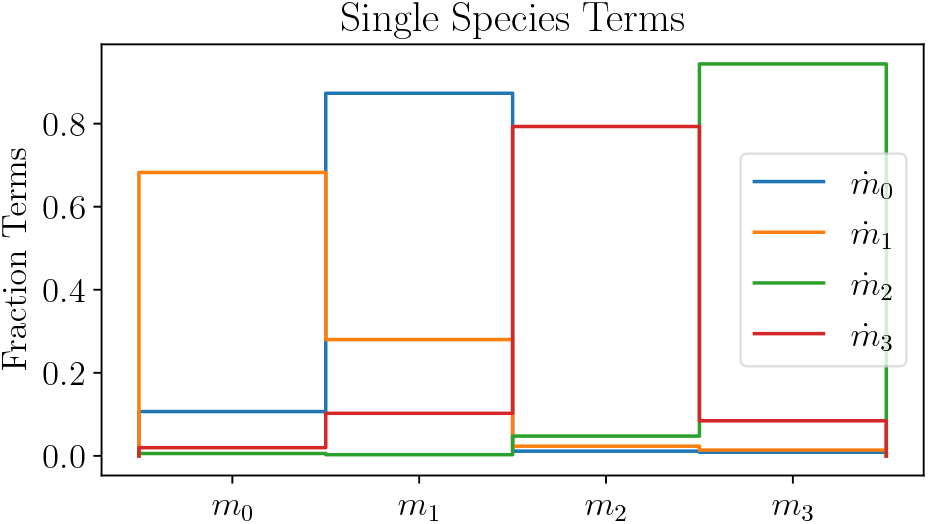
Fraction of single-species terms containing a particular species. The blue, orange, green and red lines represent 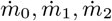 and 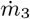, respectively. The horizontal axis represents whether the species in the single-species term is *m*_0_, *m*_1_, *m*_2_ or *m*_3_.

In Fig. 5, we plot distributions of the coefficient and thresholds for the most important single-species terms in each equation. These lines correspond with the peaks in Fig. 4. Once again, 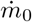 and 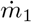 have similar properties. For 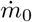, the coefficient takes the value *g*_01_ = 1.00 ± 0.42 and the threshold takes the value *K*_01_ = 0.76 ± 0.25. Meanwhile, 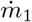 has *g*_10_ = 1.05 ± 0.54 and *K*_10_ = 0.714 ± 0.44. The main difference is that the peaks for *g*_01_ and *K*_01_ are narrower and taller, especially in the case of the threshold. The thresholds are notable for their proximity to the final stead-state values for *m*_0_ and especially *m*_1_ so that the steady state occurs when they are near their steady state. Moreover, the threshold effect is important for these species as they traverse a large fraction of the transition from low species concentrations to half of their saturation value. The coefficient for 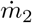 is similar to 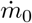 and 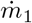 with *g*_23_ = 1.07 ±0.48, however 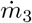 stands out sharply from the others with *g*_32_ = 0.69 ±0.44. Presumably this is because *m*_3_ needs to decrease following dilution and therefore needs to reduce this positive feedback since it would tend to cause *m*_3_ to increase. However, it should be remembered from Fig. 3 that single-species terms are only a small contribution to 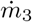. Further, their thresholds are *K*_23_ = 1.12 ± 0.36 and *K*_32_ = 1.07 ± 0.57, which are very similar, although *K*_23_ is narrower. Interestingly, the data appear to favor a threshold for *m*_2_ and *m*_3_ that are well above both their initial steady state and their final steady state. This suggests that the threshold may not be very important and they spend the majority of their time well below saturation. Therefore, the positive feedback from these terms is nearly linear. This is interesting since both have relatively little change between their diluted value and their final steady state.

**FIG. 5.**
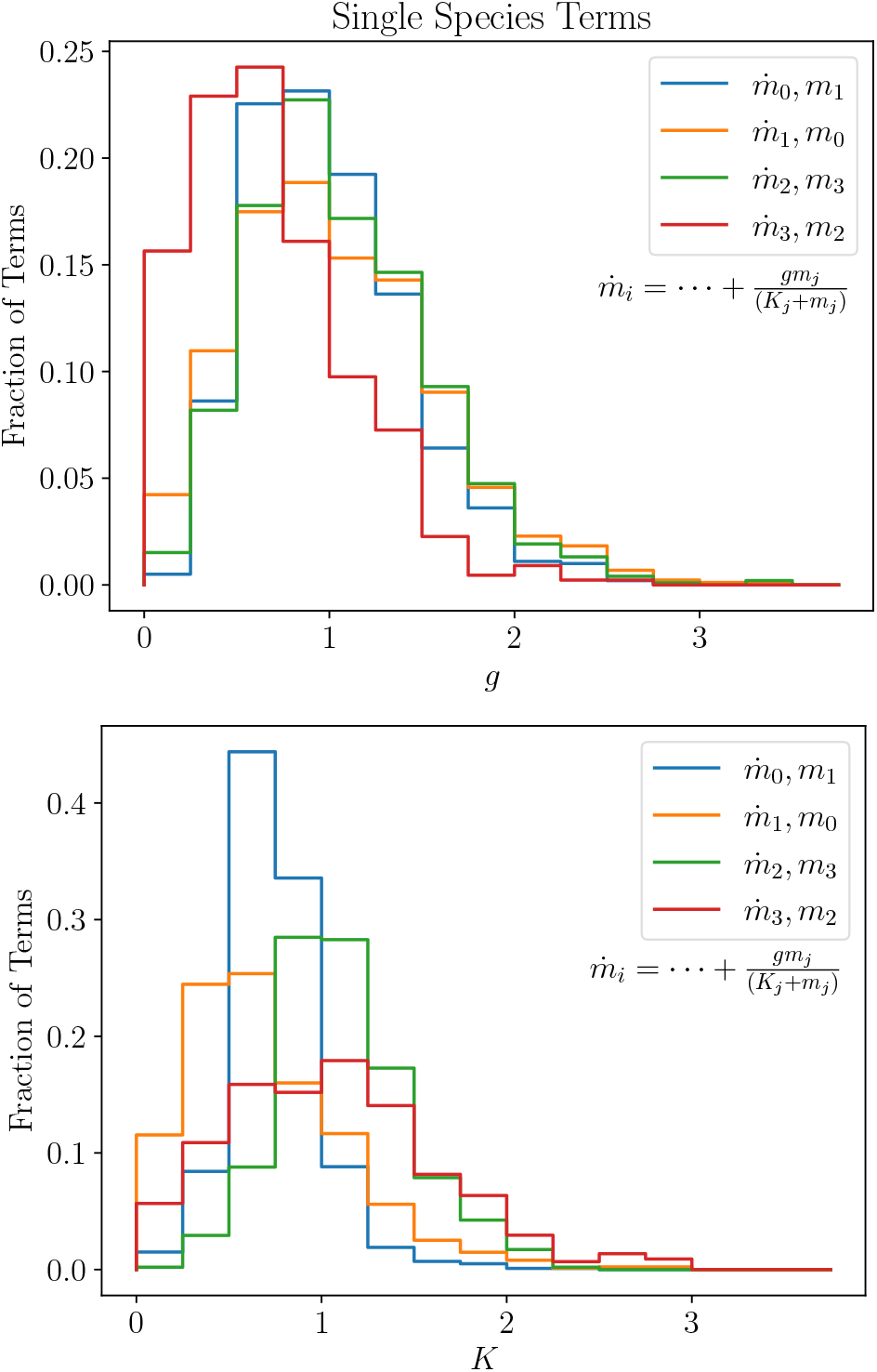
Percent of single-species terms with coefficients (top) and thresholds (bottom) with given values (horizontal axes). The coefficients and thresholds refer to *g* and *K*, respectively, in Eq. (A2). For each equation 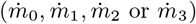, we have only shown the coefficients and thresholds for the term that occurs most frequently. That is, for the peaks in Fig. 4.

In Fig. 6, we plot which species are included for double-species terms for each equation. These are terms of the form Eq. (A4). The first thing that we note is that, in the equation for 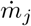 where *j* = 0, 1, 2 or 3, the majority of double-species terms contain themselves. In other words, they contain a product of the form *m*_*j*_*m*_*k*_, where *k ≠ j*. So, for example, approximately 71%, 96%, 70% and 68% of the double-species terms for 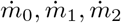 and 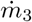 contain a product of the form *m*_0_*m*_*k*_, *m*_1_*m*_*k*_, *m*_2_*m*_*k*_ and *m*_3_*m*_*k*_, respectively, where *k* is some other species. For the equation for 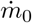, the other species in *m*_0_*m*_*k*_ is spread nearly evenly with between 20% and 30% for each of *m*_1_, *m*_2_ and *m*_3_. For 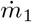, on the other hand, a dominant 60% of the double-species terms contain *m*_1_*m*_3_, where *m*_3_ is the only species that decreases following dilution, while *m*_1_ is the species that increases by the most. Additionally, 34% of the terms for 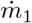 contain *m*_1_*m*_2_, where *m*_2_ is the other species whose final steady state is lower than the initial one. It appears to be important for *m*_1_, which grows substantially, to have a negative (see the next paragraph) feedback mechanism linked to the species that decrease. On the other hand, nearly no terms have *m*_1_*m*_0_, where *m*_0_ increases following dilution. Turning to 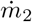, 40% of the double-species terms contain *m*_2_*m*_3_, perhaps because they share the property that their final steady state is lower than their initial one. Finally, for 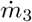, its most frequent double-species term is *m*_0_*m*_1_, which interestingly does not contain *m*_3_. Apparently, the decreasing *m*_3_ needs a positive (see the next paragraph) feedback depending on the levels of the increasing *m*_0_ and *m*_1_. However, there are nearly as many equations with a term of the form *m*_2_*m*_3_ and *m*_1_*m*_3_, both of which do contain *m*_3_. This unique feature must be due to the unique evolution of *m*_3_ following dilution.

**FIG. 6.**
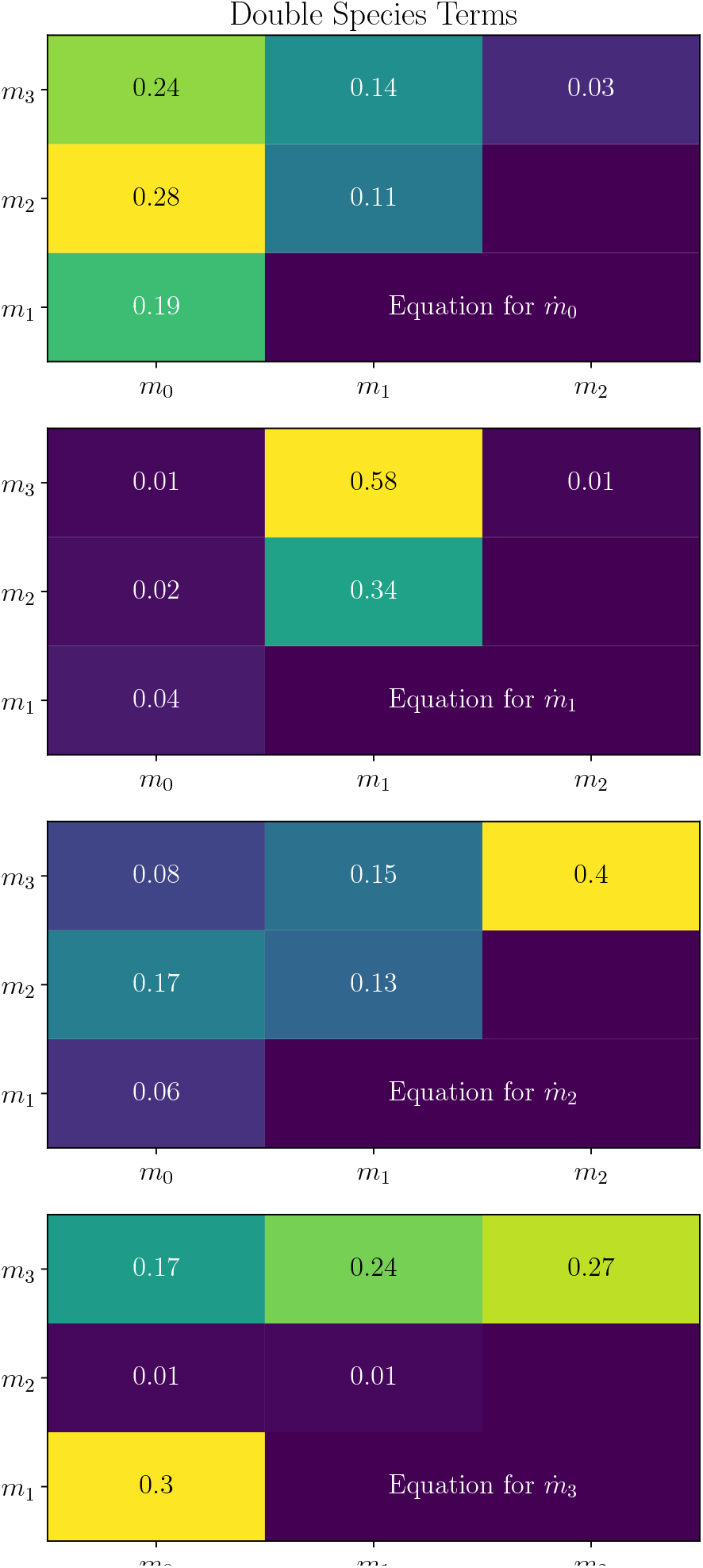
Fraction of double-species terms containing two species. The plots are, from top to bottom, for 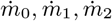 and 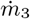. The horizontal and vertical axes are for the two species in the product.

In Fig. 7, we plot the coefficients of the most important double-species terms. As can be seen, we do not include any double-species plots for 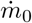 or 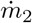. The reason is that the statistics are quite poor as can be deduced from Fig. 3 combined with Fig. 6 and therefore these terms are not present in a large fraction of models. It appears that 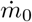 and 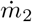 rely mainly on single-species terms. These are the two species with the smallest increase following dilution. From these same figures, we can also see that 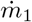 uses double-species terms much more frequently and that half of its terms contain a product of two species. Presumably this is because it increases the most drastically following dilution and needs to be closely coupled with other species in order to time both its increase as well as its approach to equilibrium. 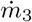 uses, by far, the greatest number of double-species terms at greater than 80% of its terms. This must be due to the fact that it is the only term that decreases following dilution. Again, we think that these terms better enable it to time its evolution to a new steady state. For these reasons, we focus on 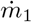 and 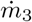.

**FIG. 7.**
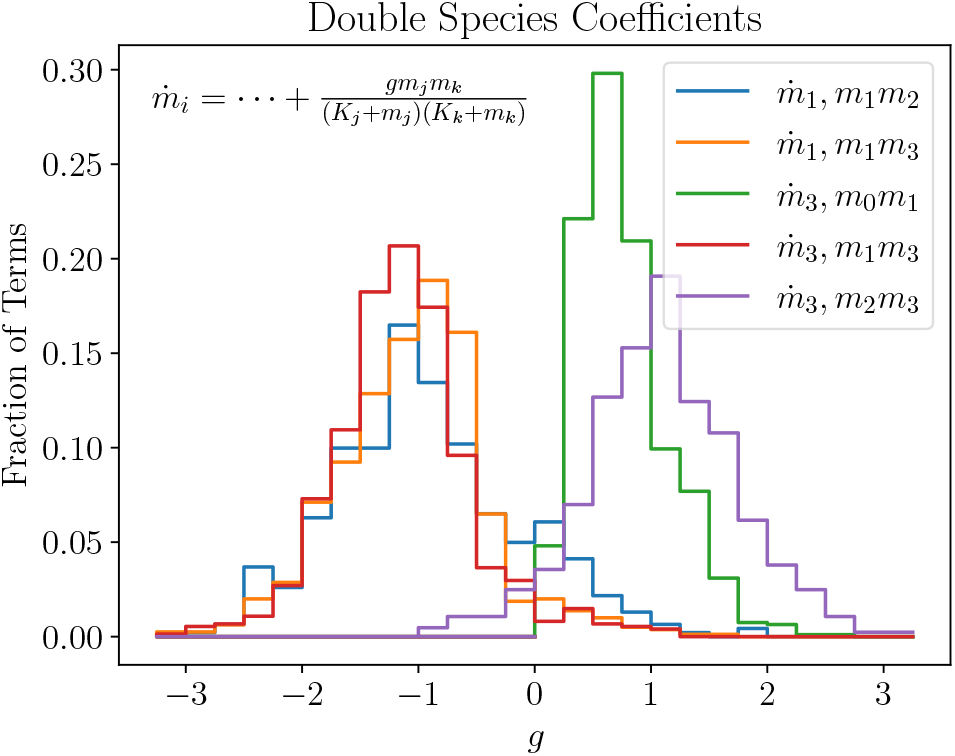
Fraction of double-species terms with coefficients of given values (horizontal axis). The thresholds for these terms are shown in Fig. 8. As described in the text, we have only shown the coefficients for the terms that occur most frequently.

As we look at the (blue and orange) distributions for 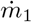 in Fig. 7, we see that both are negative. In particular, *g*_112_ = −0.91 ± 0.79 and *g*_113_ = −1.05 ± 0.65. This makes sense since, as we can see in Fig. 5, the single-species terms tend to feedback positively causing it to increase its amount. In order for this positive increase to end, 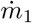 needs a negative feedback mechanism that keeps it from increasing too much. This negative feedback should be sensitive to both its own value as well as the value of other important species, especially to *m*_3_. The coefficients for 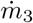 (the green, red and purple lines in Fig. 7), on the other hand, are split between positive and negative feedback. Namely, *g*_301_ = 0.76 ±0.39, *g*_313_ = −1.14 ±0.60 and *g*_323_ = 1.09 ± 0.65. For 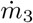, the desired effect is just the opposite. As we can see in Fig. 5, the single-species terms tend very strongly to be positive feedback and thus tend to increase the concentration of the species. For this reason, it appears, 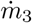 does not have an abundance of single-species terms. In fact, more than 80% of its terms are double-species terms, as noted previously. We will come back to this point in the next paragraph. In order to decrease following dilution, 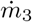 relies on joint negative feedback from itself (as is necessary) as well as *m*_1_ (the red line). Since this term tends to decrease the amount of *m*_3_ along with its decay, it now needs positive feedback to compensate and allow it to reach a steady state. It does this with the combination *m*_0_*m*_1_, which does not depend on itself and, therefore, must be positive, as well as from *m*_2_*m*_3_.

In [1], the authors noted that it was interesting that many constituents of the plasma increased rather decreased following dilution. This might be surprising since, naively, one might expect a dilution to reduce the concentrations to a lower steady state. Moreoever, inspecting their Fig. 5 reveals that all of the reported mouse proteins increased beyond their original values while nearly 80% of the reported human proteins increased. This appears to suggest that it is more likely for a protein to increase following a dilution event rather than decrease, although they only showed a fraction of the plasma proteins and it is possible there is a bias in which proteins were shown. In this paper, we have shown that, at least within the constraints of the limited data we have, single-species terms tend to increase a species concentrations, while a decrease tends to require greater contributions from double-species terms. We might hypothesize that the reason a decrease is less likely is that the evolutionary path to enzymes that were catalyzed by a single species was more efficiently explored and that enzymes requiring two species for catalysis was more difficult to achieve stochastically and therefore, less efficiently explored. Further analysis of dynamical models might shed greater light on this biological trait.

We note one last property of these coefficients before we move on. We can see that there is a preference for larger magnitude coefficients, around 1, rather than smaller ones. Although the percentage of terms with coefficients larger than 1 decreases rapidly, it should be kept in mind that every search was begun with a random value whose magnitude was between 0 and 1. The simulated anneal process and gradient method were free to search outside this range, but their initial starting point effectively reduced the distance they searched beyond unity. We attempted to increase the initial value to a magnitude that was randomly chosen between 0 and 2. What we found was a similar deficit around 0, followed by a quick growth to a maximum value around 1, a nearly flat distribution between magnitudes of 1 and 2, followed by a quick decline afterwards. There are some exceptions, such as the green line for 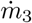 in Fig. 7 or the red line, again for 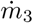, in Fig. 5, where the peaks occurs at a value lower than 1, but generally speaking, this is the behavior. These same comments apply to the coefficients of the single-species terms and the decays which were at their peak near 1 which we turn to later in this section. We suggest this to be a property of the fact that each of these models represents an infinite class of models that differ by a scaling of their coefficients and decay rates. In order to keep our analysis simple and to focus on the classes of models rather than being overwhelmed by scaled examples within a class, we chose to keep the original random range for the magnitude of our coefficients and decay rates between 0 and 1.

In Fig. 8, we show distributions of threshold values for the most frequent double-species terms. These are the *K*’s in terms of the form Eq. (A4). The main thing to note here is whether the thresholds are very small, intermediate or very large compared to the concentrations of the species during the dynamic response to dilution. We can see that there are two instances of very small threshold values and both occur for 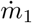 and are both thresholds for *m*_1_ (top two plots). Specifically, *t*_1121_ = 0.57 ±0.49 and *t*_1131_ = 0.48 + −0.40 (the index stands for 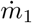 with product *m*_1_*m*_2_ or *m*_1_*m*_3_ and the threshold for species *m*_1_) both with peaks far to the left of this and long tails to the right (the mean is deceptive in this case). When this threshold is very small compared to the concentrations, it signifies that the catalyst is nearly always saturated in this species and that the reaction rate is nearly constant in this species (although not necessarily in the other species of the product). We remember from Fig. 7 that the coefficients for these terms are negative so that this represents the rate of destruction of *m*_1_. This probably could have been modeled as a single-species term without *m*_1_ in the destruction term, however our policy (see Sec. A) does not allow destructive terms unless the species being destroyed is part of the term. The reason for this was that a species has to be present in the catalyst in order for it to be destroyed and, therefore, with this constraint, the species concentration cannot become negative. The threshold for the other species in these terms has a roughly flat distribution between intermediate and large values, namely *t*_3122_ = 0.91 ±0.51 and *t*_3133_ = 0.90 ± 0.48, suggesting that saturation is not common. The dependence on the other species in this term is sensitively dependent on the concentration of the other species in some models while it is nearly linear in the other species in other models.

**FIG. 8.**
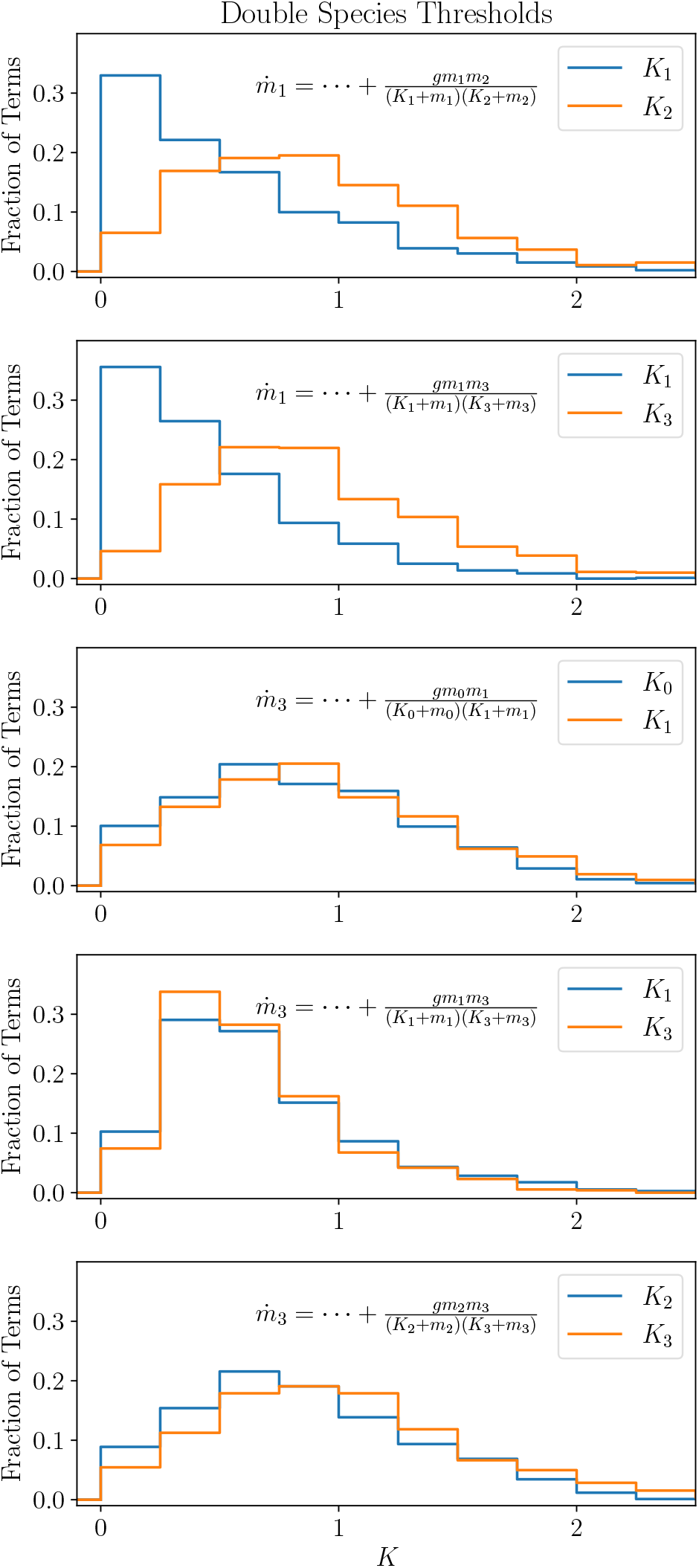
Fraction of double-species terms with thresholds of given values (horizontal axis). The coefficients for these terms are shown in Fig. 7. As described in the text, we have only shown the thresholds for the terms that occur most frequently.

Turning to 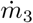, we see that only one term has a strong preference for intermediate thresholds (the fourth plot of Fig. 8). It is interesting that this is the only term with a negative coefficient (see Fig. 7) and depends on both itself *m*_3_ as well as *m*_1_. These thresholds are *t*_3131_ = 0.68±0.41 and *t*_3133_ = 0.65 ±0.36, again with a peak to the left and a long tail to the right. As we have seen in previous paragraphs, these are typically the species that stand out and this is no exception. This is the only major catalyzed destructive reaction for *m*_3_ and since the threshold value is intermediate for both *m*_3_ and *m*_1_, the rate of destruction varies greatly by the concentrations. In particular, directly following dilution, it is at its greatest rate and nearly linear. However, as *m*_1_ increases towards its final steady-state value, it approaches and then surpasses its threshold value, slowing down the rate until it is nearly constant in *m*_1_. This likely helps *m*_3_ to stabilize at its final value and not decrease too much. The dependence of the rate on itself *m*_3_, on the other hand, is nearly constant before the dilution when *m*_3_ is high, but approaches linear in *m*_3_ after the dilution and remains so to the final steady state. This makes sense since, if it increased from its steady state value, it would want to pull back at a level commensurate to its distance from the steady state. The other thresholds appear to disfavor very low values but only have slight to no favor between intermediate and larger values, namely *t*_3010_ = 0.88 ±0.51, *t*_3011_ = 0.96 ±0.54, *t*_3232_ = 0.86 ±0.49 and *t*_3233_ = 1.01 ±0.53. This suggests that a rate that is either significantly varying or nearly constant are adequate in the concentrations but a linear rate would be too much.

In Fig. 9, we consider the decay rates for each species. The *λ* refers to the coefficient in Eq. (A1). The red line for *m*_3_ immediately stands out. It has a very strong preference for very small decay rates, *λ*_3_ = 0.14 ± 0.12. This suggests that its decrease following dilution depends on the species having a relative stability compared to the other species. The destruction of *m*_3_ apparently depends heavily on catalysts and it does not tend to break apart on its own (on the scale of these experiments). The next lowest decay rate is for *m*_1_, *λ*_1_ = 0.46 ±0.27, another term which has a strong destructive contribution from catalysts and this is correlated with relative stability on its own. Although in this case, its uncatalyzed decay does happen at an appreciable rate on the time scale of the reactions. *m*_2_ and *m*_0_ have decay rates that are a bit higher, *λ*_2_ = 0.75 ±0.34 and *λ*_0_ = 1.02 ± 0.43, respectively. This is noteworthy because *m*_2_ and *m*_0_ are the two species that do not have strong catalyzed destructions. So, they make up for this by falling apart more easily, thus stabilizing the catalyzed creations of them in order to achieve a steady state.

**FIG. 9.**
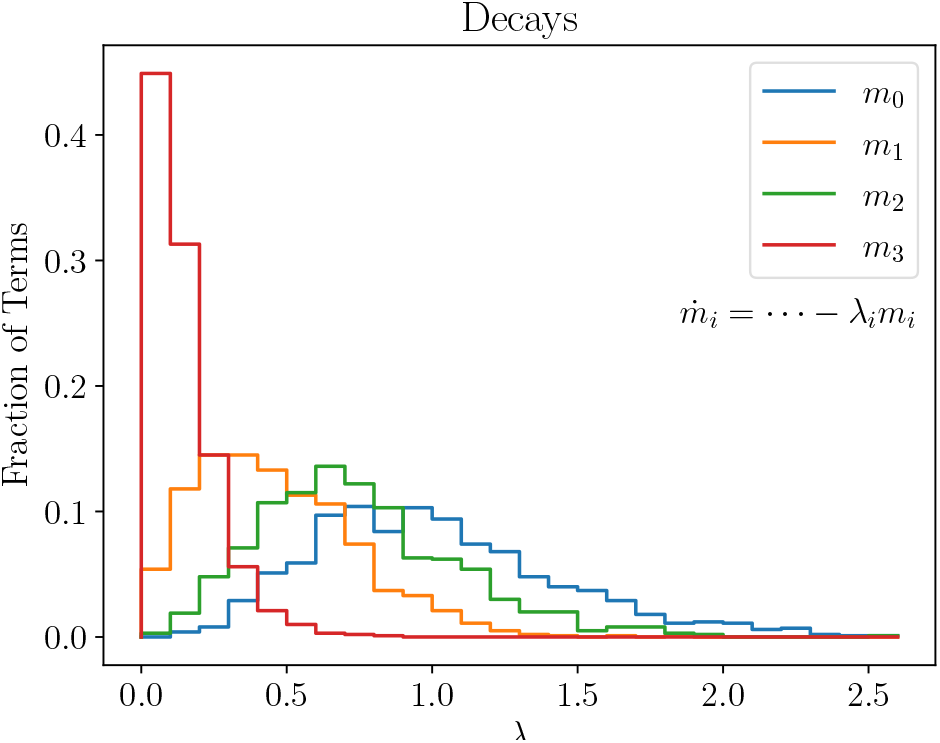
Fraction distributions of the decay rates for each species.

In this section, we consider statistical distributions of models that fit the illustrative data in Eq. (6). Since the data for these four species was so limited, there was a very large amount of ambiguity in which model could fit the data. However, we have shown that some correlations could nevertheless be inferred that might be useful as more data is collected and the models are improved. We also note that these models and the associated statistical inferences are further limited by the ability of the algorithms to generate instances of unique classes of models that fit the data. We have attempted to make a robust algorithmic system that does not prefer any type of model, but we can never be sure and further studies are always warranted. Finally, due to our limited computational resources, we were necessarily constrained to including a very small set of measured species, only 4 in this analysis, and no extra species that were not measured. This is important because the presence of feedback interactions with other species not included here, could modify the results of this study. It may be that the dependence of one species on another found here might be shifted to a dependence on other species which were not modeled here. This is a major direction we would like to consider in future studies by enlarging our computational power and analyzing models with a greater number of species. Moreover, we would like to include further data, both with a variety of dilution values as well as data at more time points. This last point, of course, will depend on the experiments being performed, but will lead to the greatest improvement in these models.

## II. DILUTION VARIATIONS

In this section, we consider variations on the dilution from the experiment. We would like to explore what predictions a dynamical system trained on sufficient data might make. Since we do not have much data for training, we cannot expect the predictions of individual models to match Nature yet. Instead, we have a population of models and, therefore, a population of predictions, each unique. On the other hand, we can see patterns among the models, suggesting that there might be a greater likelihood of a prediction to have certain characteristics. Nevertheless, any predictions made at this point are premature and should be seen as illustrating what might be achieved had we sufficient data to find a more robust, realistic model of an actual biological system. We note that all the figures in this section have dilutions that range from 0.02 up to 1.4. Dilutions above 1.0 are, of course, not dilutions. They are enhancements (infusions). However, for convenience of language, when referring to them together with the dilutions, we will simply call them all dilutions. On the other hand, when we focus on values larger than 1, we will use the more appropriate term enhancements.

We begin by considering a flat dilution where we dilute every species by the same amount, although not necessarily the 50% discussed in Sec. I. In Fig. 10, we show the range of final values for each of the species if they are all diluted by the same amount, shown along the horizontal axis. The solid curves represent the median values while the shaded regions show the range of the middle 67% of the models. We can see that if we dilute all of the species by any value below 1, 67% of the models evolve to the same final steady state that occurs with a 50% dilution. We will call this the original final steady state (OFS). This suggests that in these simple models, the OFS is an attractor, and we will see further evidence of this in the rest of this section. If we instead enhance the species, we see that the majority of the models tend towards final values where *m*_0_ and *m*_1_ are nearly 0 while *m*_2_ tends towards a range between 0.98 and 1.4 and *m*_3_ tends towards a range between 2.3 and 11.3, all in 67% of our models.

**FIG. 10.**
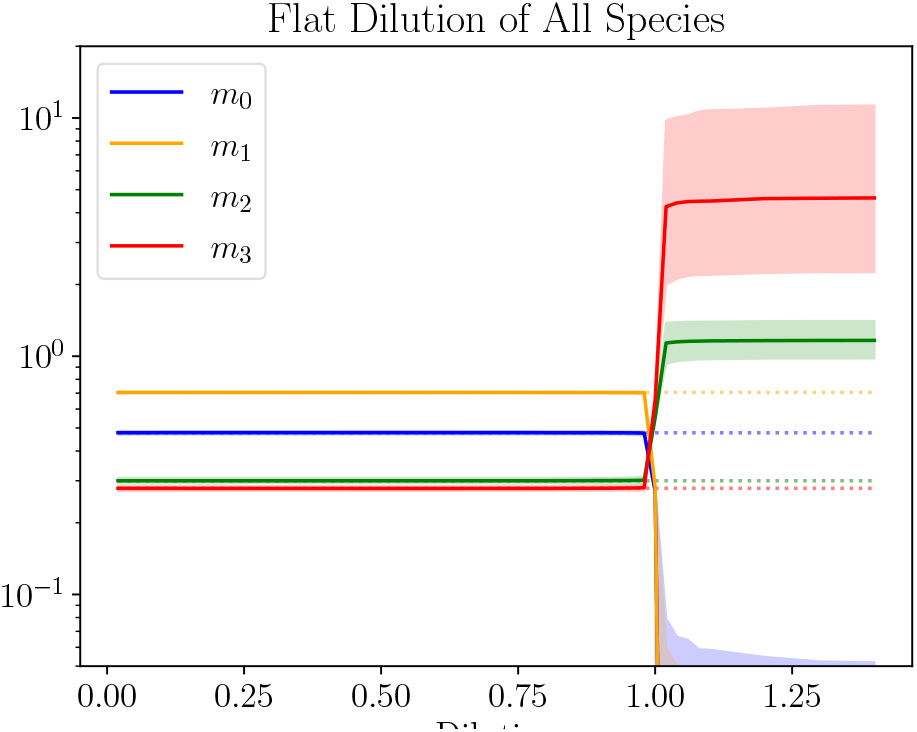
Final Values following a flat dilution. The shaded region covers the final value of 67% of the models. All the species were diluted by the same amount, shown along the horizontal axis. The dotted lines give the final steady states of the original data (the OFS).

In Fig. 11, we consider a flat 50% dilution for all the species except one. The excluded species is instead diluted at a range of values including greater dilution, lesser dilution and enhancement. If the dilution or enhancement is greater than 50% for the decoupled species, this could be achieved by a dilution followed by an infusion of the separated species. However, if the singled-out species is diluted by a greater amount than the rest of the plasma, this could not be done with a dilution-infusion combination. Rather, this might require the infusion of some agent that could bind to the species and remove it in some way. These comments also apply for other scenarios discussed in this section.

**FIG. 11.**
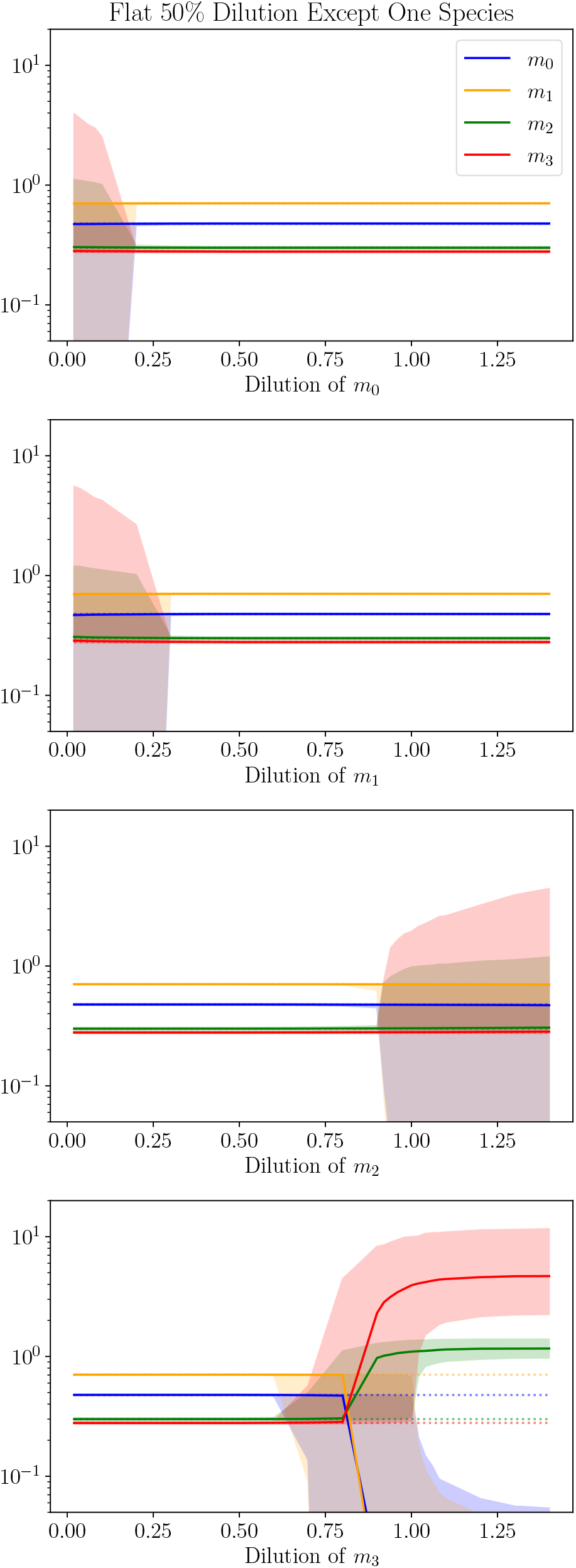
Final Values following a 50% dilution of all species except for one. The single species not diluted by 50% had a varying dilution given by the horizontal axis. The shaded region covers the final value of 67% of the models. The dotted lines give the final steady states of the original data (the OFS).

We begin with the top plot which considers a decoupled *m*_0_ dilution. We see that, given a 50% dilution for everything else, the system is very insensitive to the dilution value for *m*_0_. For nearly the entire range, the final value coincides with the original final steady state (OFS) for 67% of the models. If the dilution reaches down below approximately 0.2, the final steady states begin to spread out, although the median value still coincides with the OFS. A very similar statement can be made for dilutions of *m*_1_ and *m*_2_. For the majority of dilution values, the final steady coincides with the OFS. However, if the dilution of *m*_1_ falls below approximately 0.3 or the dilution value of *m*_2_ falls above approximately 0.9, the final steady state begins to spread out, once again with the median remaining true to the OFS. These are the three species which increase following dilution. The main difference between *m*_0_ and *m*_1_ on the one hand and *m*_2_ on the other is that they are sensitive to opposite ends of the dilution spectrum. The reason for this is that the OFS for *m*_0_ and *m*_1_ are above their initial steady state, therefore they are more sensitive to severe dilutions which are in the opposite direction of their OFS. The behavior of *m*_2_ is the opposite with its OFS below its initial steady state, therefore it is more sensitive to enhancements which lead in the opposite direction of its OFS.

The species that really stands out here is *m*_3_, which is sensitive to both dilutions above 0.7 as well as enhancements. Or to put it differently, in order to reach the final steady state of the data, *m*_3_ must be diluted by at least 30%. If it is above that, even with all other species diluted by 50%, the model will tend towards a different final steady state. *m*_3_ is the most unusual as we have seen throughout this note as it, not only has a lower final steady state, but it also decreases following dilution and requires the greatest complexity of its dynamical equations. It appears that this confers a greater sensitivity on it than the other species. Interestingly, with enhancements, 67% of the models tend towards final values with *m*_0_ and *m*_1_ at or very near 0 while *m*_2_ and *m*_3_ tend towards values in the range 0.97 and 1.4 and 2.3 and 11.6, respectively. These are the same final values as occurred in Fig. 10, where all species were enhanced by the same amount. This suggests that *m*_3_ might have been the dominant change causing evolution towards this final state. Having the other species diluted at 50% here helped the transition occur so that *m*_3_ could take a larger range of dilutions and enhancements of 0.7 and above to achieve these final states compared with the flat dilutions shown in Fig. 10.

We also considered manipulation of two species independently while keeping the remaining two at a 50% dilution. So, for example, we diluted *m*_2_ and *m*_3_ at 50% and then varied the dilution or enhancement of *m*_0_ and *m*_1_, independently, over the range 0.05 to 1.4. We found roughly what we would expect if we considered the combined effect of two modified dilutions that are shown in Fig. 11. Since there are too many plots to display here, and since there is a large degree of similarity between the plots, we will focus on a few illustrative examples here. In Fig. 12, we present the median final value of *m*_1_. Along the horizontal axes and vertical axes, we present the dilution values for two species while we hold the dilution of the other two species at a constant 50% dilution. So, although the final value after dilution is always presented for *m*_1_, the dilution values depend on the values of the axes or are 50% if not on the axes. In these plots, the yellow region shows where the median final value of *m*_1_ is the original final steady state (OFS), the right side of Eq. (6). The blue-green area is the transition region between final states and the dark blue region shows the region where the alternate final steady state (AFS) occurs following dilution, shown on the right side of the bottom plot of Fig. 11, for example. Although we only show plots for the final state of *m*_1_, the plots for the other species, given the same dilutions, are analogous. The exact positions of the the lines separating the regions is slightly altered but is in the same region and has the same qualitative shape.

**FIG. 12.**
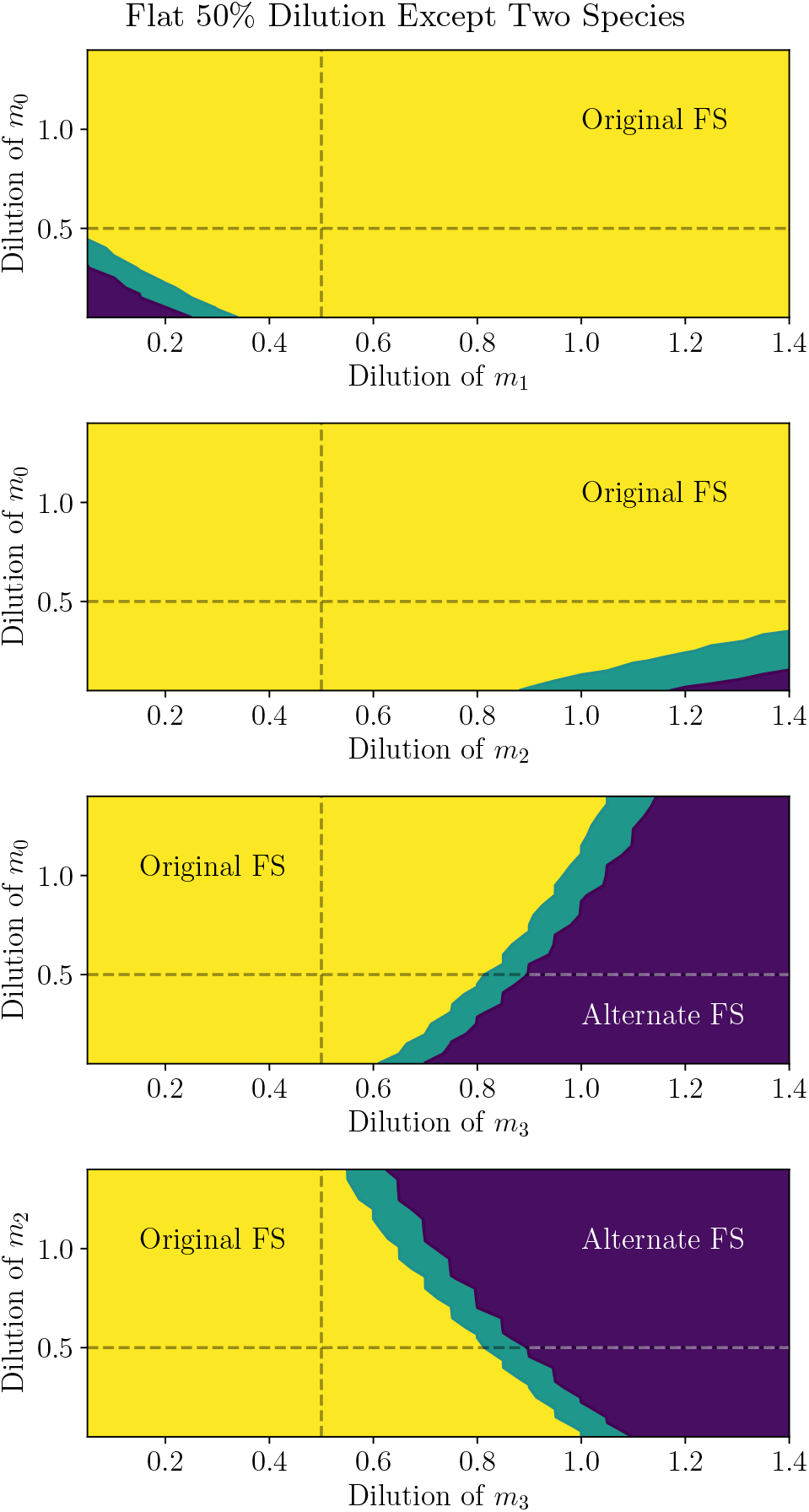
Distributions of median final value of *m*_1_ when two species were diluted or enhanced, independently. These two species are specified in the horizontal and vertical axes. The other two species, not on the axes, are diluted at 50% in each case. FS stands for final steady state.

In the case that we dilute *m*_2_ and *m*_3_ at a constant 50% but vary the dilution of *m*_0_ and *m*_1_ (top plot of Fig. 12), we find that, for the majority of dilution and enhancement values, the final state is the OFS. However, if we dilute either *m*_0_ or *m*_1_ to very small dilutions, the final state switches to the AFS. This is shown as the darker region at the bottom left. This is because, as we can see in the second plot of Fig. 11, the final state begins switching at very low dilutions of *m*_0_ and *m*_1_ and when we combine very low dilutions of both, the switch becomes complete. We note that the solid orange line in the second plot of Fig. 11, corresponds with a horizontal line in this plot at a height of 0.5, and is yellow for the entire length, as expected. Similarly, the solid orange line in the first plot of Fig. 11, corresponds with a vertical line in this plot at a value of 0.5, and is also yellow for the entire height, as expected. Similar comments apply for the horizontal and vertical dashed lines in the other plots of Fig. 12. We also note that we are only plotting the median value and don’t have a clear way to include the 67% confidence region. In the second plot of Fig. 12, we show the effect of varying the dilution of *m*_0_ and *m*_2_. We can see that the alternate final steady state switches from the bottom left to the bottom right. This makes sense since, as we can see in the third plots of Fig. 11, the final state switches for high values of *m*_2_, which is the right side in this plot. Again, we find that low values of *m*_0_ hasten the transition. If we vary the dilutions of *m*_1_ and *m*_2_ instead (not shown), we find a qualitatively very similar plot to the second plot of Fig. 12. Again, the transition region is at the bottom right. However, the transition line has a slope that is shallower.

The shape changes significantly when we vary the dilution of *m*_3_, as we would expect based on the bottom plot of Fig. 11. If we also vary *m*_0_, we get the behavior shown in the third plot of Fig. 12. The transition is most sensitive to variations in the dilution of *m*_3_. However a smaller dilution value for *m*_0_ makes it more likely, while a larger dilution value (or enhancement) of *m*_0_ makes the transition less likely, causing the transition line to slope to the right. Varying the dilution of *m*_1_ along with *m*_3_ (not shown) creates a qualitatively similar plot with the slope of the transition line to the right, but the slope is shallower. Finally, if we vary the dilution of *m*_2_ along with *m*_3_, as expected, the slope of the transition line flips to the left. We expect this, based on the third plot of Fig. 11, since the transition is sensitive to large values of *m*_2_, and therefore should occur more quickly the higher the dilution value of *m*_2_.

In Fig. 13, we consider what would happen if we only diluted (or enhanced) one species but left all the others at their original value. We can see that the behavior splits into two groups. In the first, as shown in the first two plots, if we enhance either *m*_0_ or *m*_1_ alone, the model evolves towards the original final steady state (OFS). This is likely because the original final steady state is higher than the initial steady state for *m*_0_ and *m*_1_ and so this begins the evolution in the direction of the OFS. On the other hand, *m*_2_ and *m*_3_ are just the reverse. Their original final steady state is below the initial steady state, therefore, a dilution moves them in the direction of the OFS. As we see in the bottom two plots, any dilution of either of these causes an evolution to the OFS. On the other hand, if we perturb any single constituent of the group in the direction opposite the original final steady state, it evolves towards a different final state. Interestingly, that final state is essentially the same for any single-species perturbation and is, in fact, the same as when we enhance all four species (as shown in Fig. 10) or when we dilute all species at 50% except for *m*_3_, which is enhanced (as seen in the bottom of Fig. 11). Namely, in all four cases here, both *m*_0_ and *m*_1_ tend towards very small or zero values while *m*_2_ tends towards a range of 1.0 and 1.4 and *m*_3_ tends towards a range of 1.8 and 10.5 in the first plot, 2.2 and 11.5 in the 2nd, 2.2 and 10.4 in the third and 2.3 and 11.4 in the fourth. Although the range for *m*_3_ is not exactly the same, the regions overlap significantly. This alternate final steady state (AFS) region is interesting because it was not included in the training data and we did not build it into the models. It appears that, within the limitations of the data that we used, the presence of the steady states given in Eq. (6) and the evolution between the two is sufficient to predispose the model to have another steady state in this region. At least, it occurred in 67% of the models obtained using our algorithm. That is, the other 33% of our models that fit the data do not have a steady state in this region. So, if we had more data, we might continue to find a steady state in this region, or we might not. However, a more robust model, generated with a greater quantity of quality data, might predict the presence of other steady states that might be interesting and might suggest ways of achieving an evolution to the alternate steady state.

**FIG. 13.**
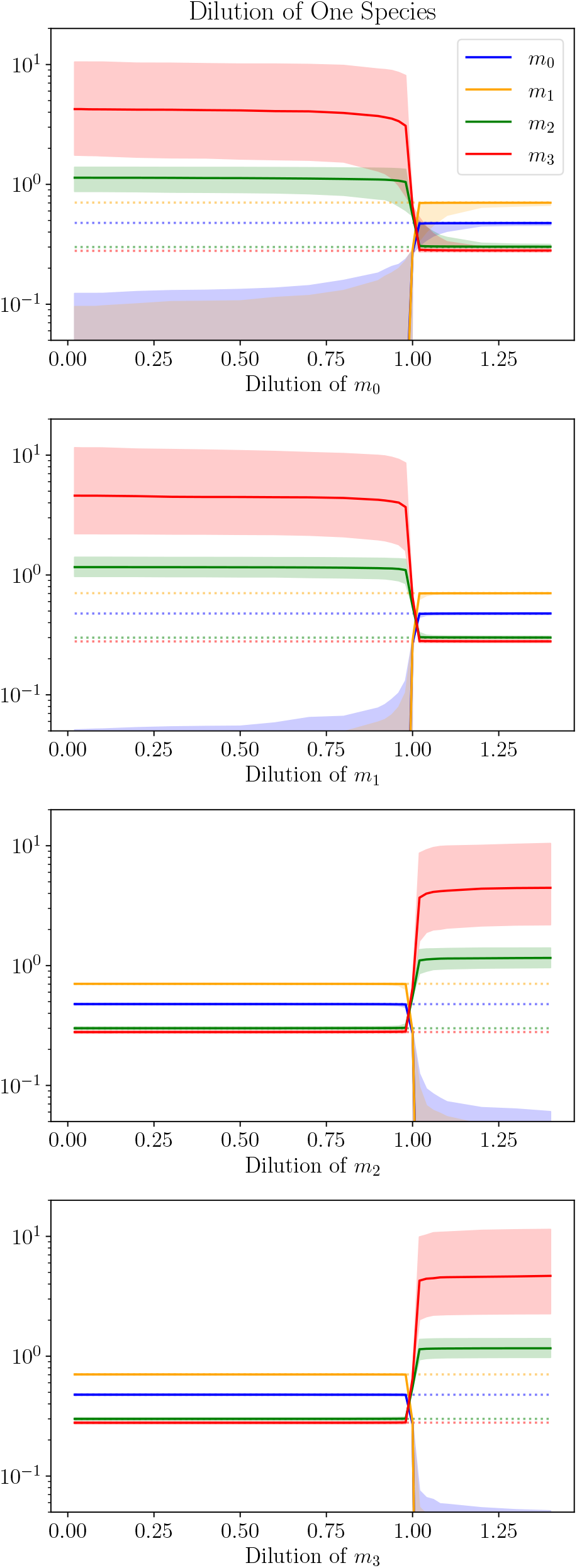
Final Values following a dilution of only one species while not diluting the others. The single species diluted is given by the horizontal axis. The shaded region covers the final value of 67% of the models. The dotted lines give the final steady states of the original data (the OFS).

Another important feature of Fig. 13 is that the original steady state is not stable. A perturbation in any direction of any single species is sufficient to drive the model away from the original steady state towards either the original final steady state or towards some other final state, as described in the previous paragraph. This property is likely not physical as we expect the levels of constituents in our plasma to be stable to small changes. This is a feature of our models we would like to improve in the future. However, we would prefer to not build in a region of stability by hand since we do not know the sizes of these regions physically and they likely are different for each species being perturbed. Furthermore, we suspect that the number of steady states as well as their stability is impacted by the size of the dynamical model. That is, as we increase the number of species in our model, including unmeasured, hidden and conserved species, as described in App. A, the number of steady states will likely increase and their stability structure could change. Moreover, we would prefer to let the data determine the features of the model. Given sufficient data of the right type, the steady states should take on more realistic properties. In particular, as a place to start, we would like to see dilution experiments with a range of dilutions, including dilutions where the plasma returns to its original steady state. The original final steady state, as given on the right of Eq. (6), on the other hand, is stable to a wide range of perturbations of a single species in our models.

In Fig. 14, we show the result of diluting two species independently, while holding the others at their original value. As in Fig. 12, we have only plotted the median final value of the models with each dilution. Moreover, as in that previous figure, we see that the dilution space is divided into three regions: the original final steady state (OFS, in yellow on the left and dark blue on the right) and the alternate final steady state (AFS, in dark blue on the left and green on the right) and an intermediate transition region separating them. However, this time there is a qualitative difference between the behavior of *m*_0_ and *m*_1_ on the one hand and *m*_2_ and *m*_3_ on the other. In particular, the transition region is broadened for the latter two. For this reason, we have plotted the distribution for *m*_1_ in the left column and these plots are also representative of *m*_0_, while we have plotted the distribution of *m*_2_ (also representative of *m*_3_) in the right column. We have also included dashed lines when the dilution of either constituent is 1.0, representing no dilution at all. These dashed lines correspond with the solid lines in Fig. 13. As expected these dashed lines cross in the transition region, where none of the species are diluted and the final state is the original starting state [the left side of Eq. (6)], which is in between the original final steady state and the alternate final steady state.

**FIG. 14.**
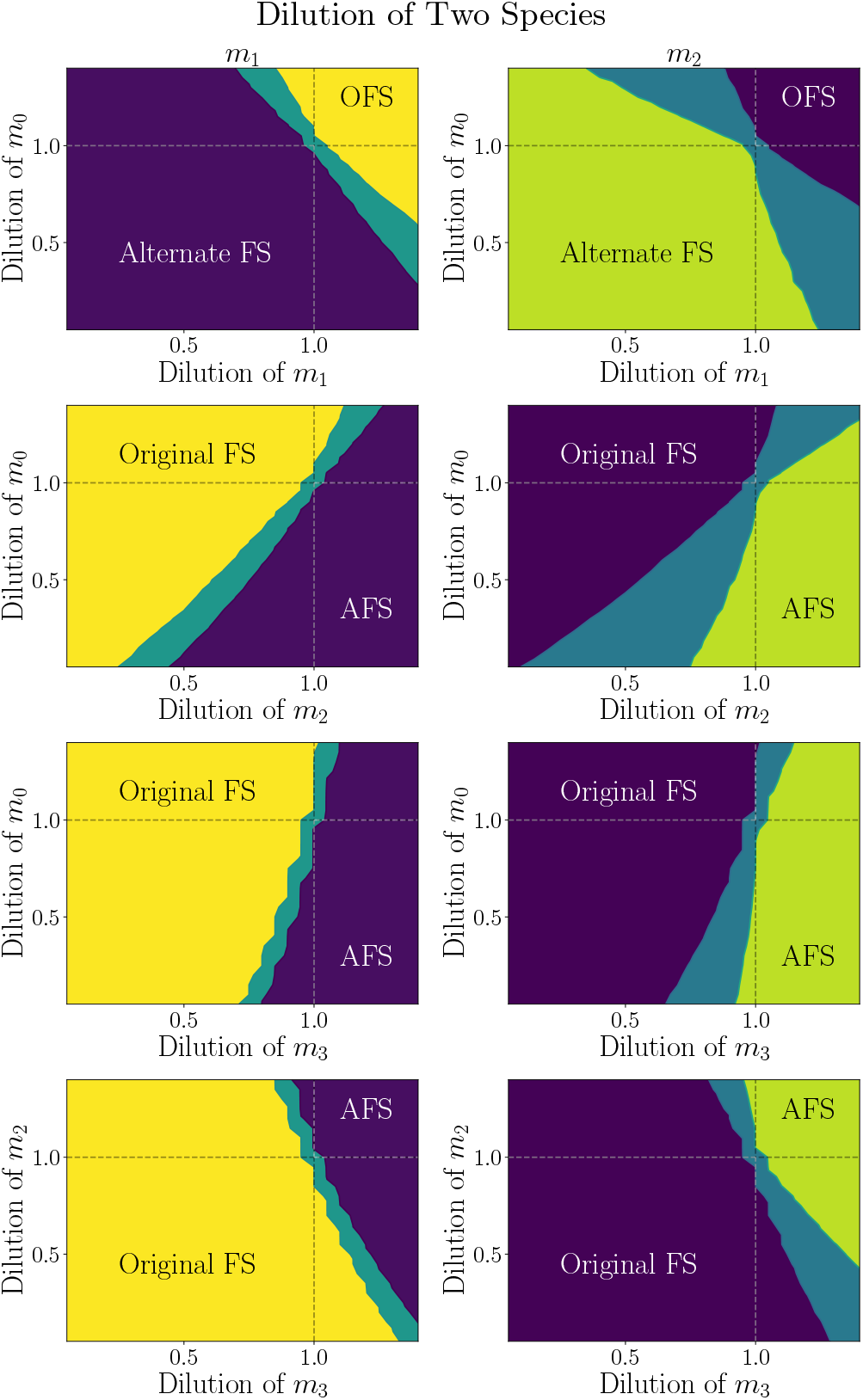
Distributions of median final value of *m*_1_ on the left and *m*_2_ on the right when two species were diluted or enhanced, independently. These two species are specified in the horizontal and vertical axes. The other two species, not on the axes, are held at their original value in each case. FS stands for final steady state.

In the top row of Fig. 14, we see the behavior of the constituents when only *m*_0_ and *m*_1_ are diluted. As we would expect, based on first and second plots of Fig. 13, the original final steady state is obtained if both species are increased, but the alternate final steady state is preferred if they are both diluted. Additionally, in the top left and bottom right regions, we see that an enhance-ment of one but a dilution of the other results in the OFS if the dilution is not too much, but transitions to the AFS if the dilution is large. We also see, in the right plot for *m*_2_ and *m*_3_, that the transition region takes up a larger fraction of the space in the top left and bottom right, when one species is enhanced and the other diluted. This is presumably because *m*_2_ and *m*_3_ are more sensitive to the level of dilutions/enhancements, which appears to be a consequence of their decrease following dilution. In the second row, we show plots for dilutions in *m*_0_ and *m*_2_. Once again, if we compare this behavior with the first and third plots of Fig. 13, we see that enhancing *m*_0_ and diluting *m*_2_ results in the OFS, while the opposite results in the AFS, as expected. On the other hand, either enhancing both or diluting both results in one or the other depending on which is stronger, with a transition region separating them. Once again, the transition region is larger for *m*_2_ and *m*_3_ (in the right plot). The plots (and behavior) are very similar for dilutions of *m*_1_ and *m*_2_, and so we do not include plots for this combinations here. However, we do note that the transition region is closer to the diagonal for the *m*_1_ −*m*_2_ combination. In the third row, we consider the combined dilutions of *m*_0_ and *m*_3_. In this case, the dilutions and enhancements of *m*_3_ are overpowering and the transition region is nearly a vertical line at a dilution of 1.0 for *m*_3_. We have seen throughout this note that the model is very sensitive to *m*_3_ and that this appears to be connected to the fact that, not only does it decrease following dilution, but it also decreases from its diluted value, at least with the original dilution values. Nevertheless, we do see a little slant to the transition line showing that enhancments of *m*_0_ do increase the OFS region a little and dilutions of *m*_0_ decrease it a bit, as expected. The combined *m*_1_− *m*_3_ dilution behavior is very similar to this and we do not show it. On the last row, we consider dilutions of *m*_2_ and *m*_3_. Since dilutions and enhancements tend to have similar effects, we see that dilutions of both give the OFS while enhancements of both give the AFS. Enhancement of one and dilution of the other is split, depending on which is stronger. This case is the opposite of the first row, where we consider the *m*_0_ − *m*_1_ case.

## III. SUMMARY AND CONCLUSIONS

In this note, we considered dynamical systems models for plasma dilutions. The plasma dilution experiments, so far, have only been done at one dilution value and the concentrations of plasma proteins have only been measured once before and once after the dilution event. Because this data is so limited, we were not able to achieve a unique model that faithfully represents the behavior of the plasma system in Nature beyond the measured data. That is, the predictive power of our models are admittedly and expectedly very low. Alternatively, our goal has been to show what might be achieved with a dynamical systems model had we sufficient data to properly disambiguate the models that represent the plasma system. We did this by creating a large population of models that fit the limited data and creating and analyzing statistical distributions of their properties and their predictions. Admittedly, a large number of models giving a similar prediction does not imply that Nature is necessarily more likely to satisfy the prediction. The addition of new data, especially with alternate dilution values and frequencies of plasma measurements, could, in fact, greatly alter its predictions. However, it is nonetheless suggestive that a more constrained model, given superior data, might give interesting predictions that could guide further experiments. In particular, with such a more constrained model in hand, predictions could be made based on alternate dilution values and dilution frequencies, as well as infusions. These predictions could be followed by laboratory experiments and the results of these experiments could then be fed back into the model to improve it and its future predictions. Moreover, using such a constrained and improved model to guide laboratory experiments could make the search for the optimal laboratory and clinical procedure more efficient.

In order to achieve this goal, as described in App. A, we began by developing the structure of the differential equations that would have the representational power for the plasma constituents. We considered simple decays that represented instability in the proteins in Eq. (A1) and did not require a catalyst. We also included catalyzed reactions that either created or destroyed a plasma constituent. Since catalyzed reactions were sensitive to the concentrations of each molecule attached to it, we used a model of the form Eq. (A2) for a single-species catalyzed reaction and Eq. (A4) for a reaction catalyzed by two species. With this term, the rate of production or destruction is nearly linear when the species concentrations is low compared to its threshold and nearly constant when the concentration is high compared to its threshold, and the catalyst is saturated. Additionally, we reasoned that destruction of a plasma species could only occur when the species was present in the catalyst. Therefore, we only allowed a destructive catalyst, where the coefficient is negative, when the species being destroyed was included in the catalyzed reactions. In other words, we only allowed negative coefficients in terms such as in Eqs. (A3) and (A5). This removed the possibility that the species concentrations ever became negative. All these dynamical terms were then added together to create a system of differential equations of the form Eq. (A6). We also considered different classes of plasma constituents (which we call *u*_*i*_, *h*_*i*_ and *c*_*i*_) which might be important to the dynamical reactions creating and destroying each measured species. However, only the measured species ended up playing a role in our final population of models.

In App. B, we described the computational methods that we used to create a population of models that fit the data. We used a combination of a genetic algorithm, simulated annealing and a gradient descent method. For our genetic algorithm, we began by generating a population of random models. We then trained these models using simulating annealing and gradient descent obtaining a model score that represented how close the model was to fitting the data, as seen in Eq. B2. After this, new models were iteratively created based on this population. At times these models were a combination of two models in the population, representing sexual reproduction, at other times, a new model was a “mutation” of a population model with a new term or a new species in the dynamical equations, or a term or a non-measured species removed, and at times a brand new random model was created from scratch. With each new model, it was trained, using simulated annealing and gradient descent, and its model score was compared with that of the population. If its score was better than the worst in the population, it replaced the worst model-score model. This process was continued until the model scores of the population were satisfactorily low. Simulated annealing was accomplished by randomly choosing parameter values in a hypercube around the current point, where the size of the hypercube decreased over time. To accomplish the gradient descent method, we calculated the slope at each point and moved the parameters iteratively down the slope. The gradient descent method was done for each iteration of the simulated annealing process and this seemed to give us the most efficient and successful results.

With the form of our differential equations and our algorithms in place, we ran our program for 1500 cpu-days to produce of population of 1000 models as presented in Sec. I. The model scores all fell below 0.03 as compared with the illustrative data shown in Eq. (6) and as shown in Fig. 2. Using our present computational resources, we were unable to generate sufficiently large populations of models with sufficiently low scores if we increased the number of measured species much beyond this and four species was a compromise. However, in the future, with increased access to computer clusters and with improved algorithms, it would likely be possible to scale this up to more interesting collections of data. In Fig. 3, for each species, we showed distributions of the number of terms, the number of negative terms and the number of species per term in each differential equation. From this initial point and throughout, we saw that there was a large difference between *m*_0_ and *m*_2_ on the one hand and *m*_1_ and especially *m*_3_ on the other. We noted that these were the two species with the largest change to their concentrations following dilution with *m*_3_ standing out even further because of its great reduction. In Fig. 4, we presented the fraction of single-species terms for each constituent [see Eqs. (A2) and (A3)]. In this case, we saw a strong connection between *m*_0_ and *m*_1_ on the one hand and *m*_2_ and *m*_3_ on the other. We pointed out that this was likely due to the fact that both *m*_0_ and *m*_1_ increased between their initial steady state to the their final steady state, while the latter two decreased. The single-species reactions appear to mainly be of positive feedback loops. We displayed the coefficients and thresholds of the dominant single-species terms in Fig. 5. In this case, we found that all the coefficients were positive and constructive and that the coefficient for the dominant term in the equation for 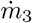 was smaller than the others and suggested this was because it not only fell from its initial steady state, but it fell from its diluted value, requiring less of this positive feedback.

We presented the fraction of two-species terms [see Eqs. (A4) and (A5)] for each equation in Fig. 6. We noted that only five of these were present in large enough numbers in the population to make their analysis interesting. The first two were for the terms with *m*_1_*m*_2_ and *m*_1_*m*_3_ in the equation for 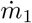. Both of these terms were negative, as seen in Fig. 7, and represented destructive feedback, reducing *m*_1_ in relation to the concentrations of *m*_2_ and *m*_3_ as well as itself. We noted that the reasons these stood out relative to other two-species terms was that *m*_1_ increased by the greatest amount and therefore it needed these negative feedback mechanisms to stabilize it at the new higher steady state against the strong positive feedback found in the single-species term. The equation for 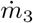, on the other hand, was mostly the reverse of this. There were three terms, namely *m*_0_*m*_1_, *m*_1_*m*_3_ and *m*_2_*m*_3_. The second of these was a negative feedback term, while the other two were positive (see Fig. 7). It appears that the challenge of decreasing to a lower steady state required more complicated dynamics. We have already seen evidence of this in the discussion of Fig. 3, where *m*_3_ required greater number of terms, greater number of two-species terms and greater number of negative feedback terms. Here, we see that it needed two-species terms that are both constructive and destructive and that achieving its dynamics is apparently a greater balance between two-species catalysts. We further pointed out that the negative feedback terms were predominantly in two-species terms, not single-species terms. We discussed that perhaps this was because negative feedback would depend on both another species concentration as well as its own. We pointed out that this might be why increased levels of protein concentrations appeared to be more likely in the results of the dilution experiments [1, 2, 9]. Perhaps concentrations are more likely to be enhanced following dilution since reductions depend more heavily on two-species catalyzed reactions and perhaps two-species catalysts are less efficiently explored than single-species catalysts by a stochastic evolutionary process.

In Fig. 9, we exhibit the decay rates for each species and find *m*_0_ and *m*_2_ are comparable while *m*_1_ is smaller and *m*_3_ even smaller. Once again, we suggested that this was because *m*_1_ and *m*_3_, with their larger changes following dilution, required a greater amount of their dynamics to come from balancing creative and destructive catalyzed reactions and less from breaking apart uncatalyzed.

In App. C, we showed a variety of examples of models from the population. We found that the population was organized into three broad groups and we presented plots and the full differential equations for four models from each group. The first group were composed of models where *m*_0_, *m*_1_, *m*_2_ began increasing immediately following dilution while *m*_3_ immediately began decreasing. Plots for exemplary models were shown in Fig. 15 while their differential equations were shown in Eqs. (C1) through (C4). The second group contained models where all of *m*_0_, *m*_1_, *m*_2_ *and m*_3_ began increasing immediately following dilution. We presented plots for illustrative models in Fig. 16 and their accompanying differential equations in Eqs. (C5) through (C8). The last group was the same as the first group except that the concentrations oscillated up and down before approaching their final steady states. Example models were presented in Fig. 17 and their differential equations were given in Eqs. (C9) through (C12).

**FIG. 15.**
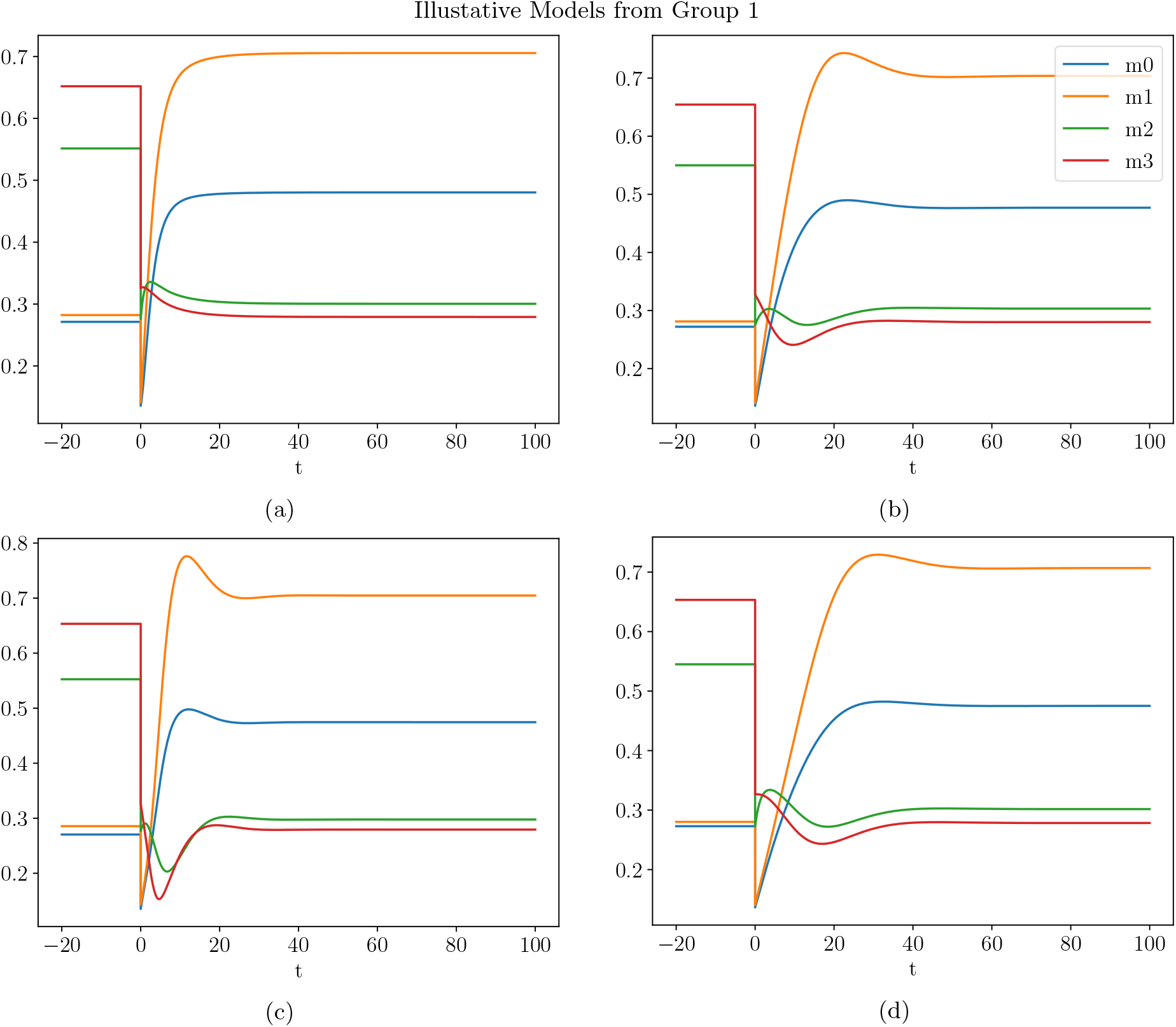
A sampling of model dilution plots from Sec. I from Class 1 (see the beginning of this appendix). The system of differential equations for these plots can be found in: (a) Eq. (C1), (b) Eq. (C2), (c) Eq. (C3) and (d) Eq. (C4).

**FIG. 16.**
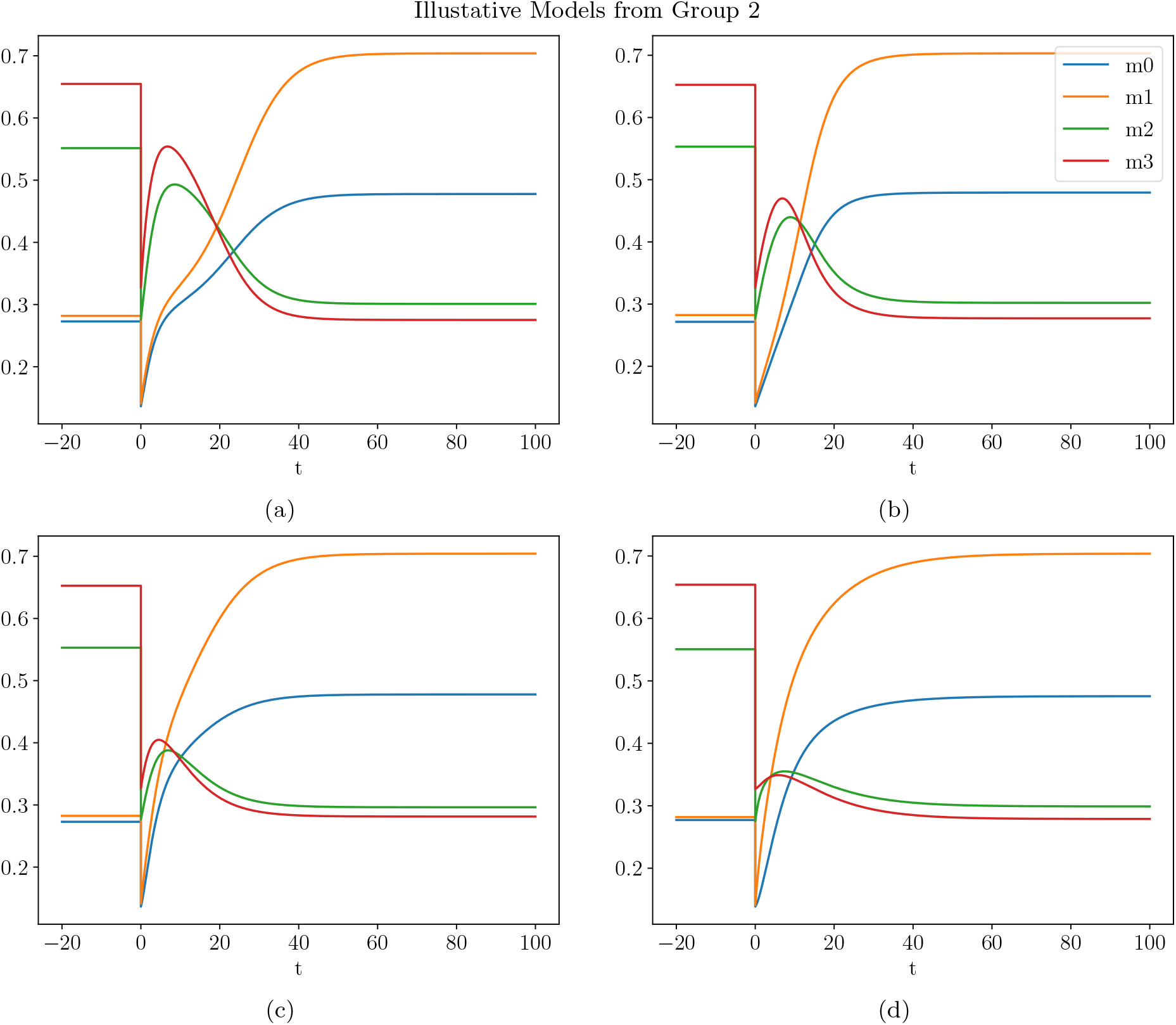
A sampling of model dilution plots from Sec. I. The system of differential equations for these plots can be found in: (a) Eq. (C5), (b) Eq. (C6), (c) Eq. (C7) and (d) Eq. (C8).

**FIG. 17.**
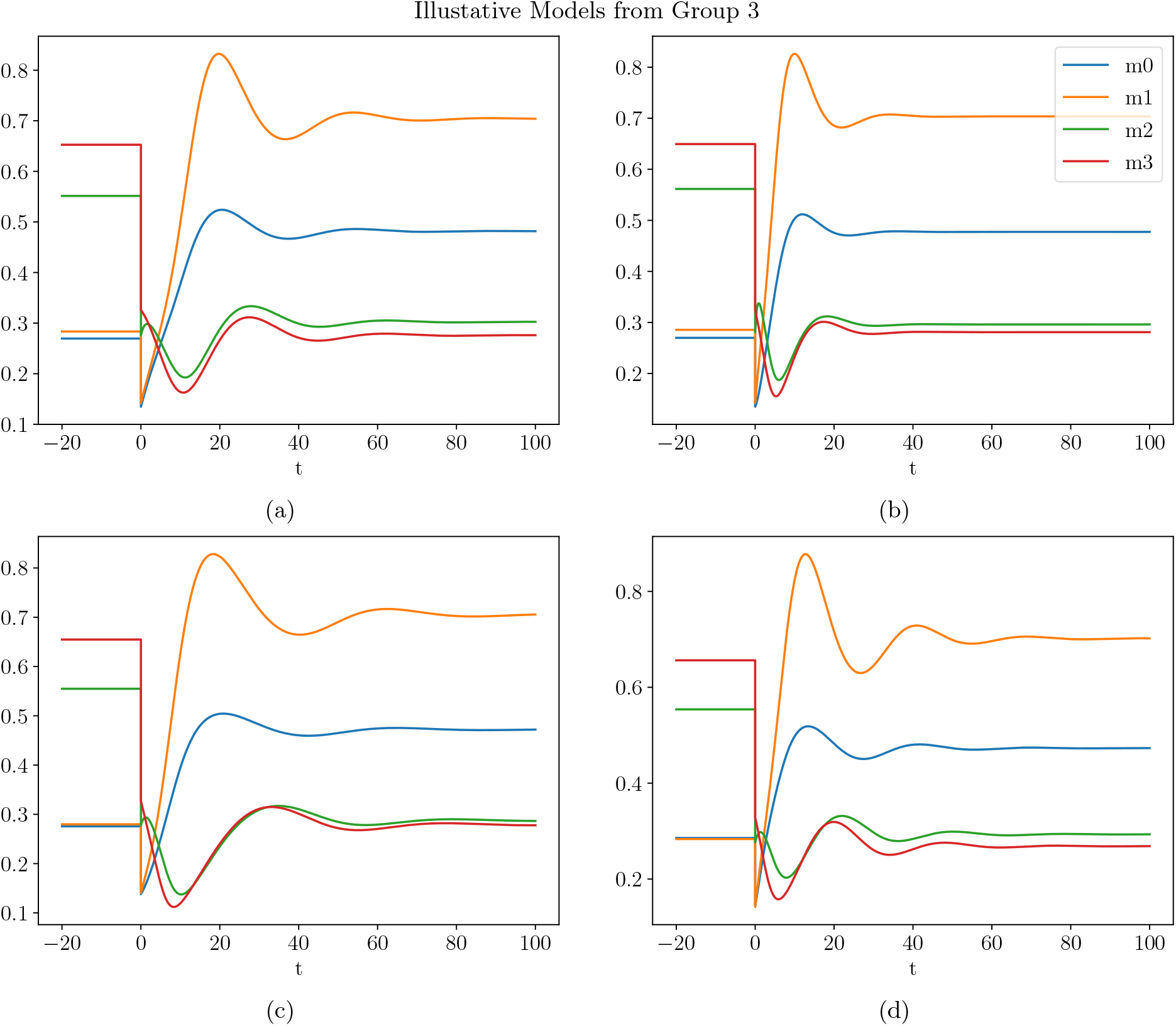
A sampling of model dilution plots from Sec. I. The system of differential equations for these plots can be found in: (a) Eq. (C9), (b) Eq. (C10), (c) Eq. (C11) and (d) Eq. (C12).

In Sec. II, we explored and illustrated the properties we might learn about the dynamical system and about alternate dilution values had we greater data and a more realistic, more robust model. In Fig. 10, we considered the behavior if we diluted all constituents by a flat amount but at different dilution values. We found that for any dilution values, the system was likely to end up in the original final steady state (OFS), but that if, instead of a dilution, an infusion of all four species were achieved, the system would evolve to an alternate final steady state (AFS). Although it isn’t likely we would infuse all the constituents of plasma in this manner, the AFS in this plot showed up in every other case we discussed in this section. In Fig. 11, we considered what would happen if we diluted all species at the original flat 50% except for one species which was diluted at a variety of levels. If the single species diluted at a different level were one of *m*_0_, *m*_1_ or *m*_2_, then the system would typically still evolve to the OFS, even if the single species was instead enhanced. However, the model proved especially sensitive to the dilution value of *m*_3_. Indeed, we found that any dilution of *m*_3_ above approximately 70% would lead the system to finish at the AFS. Interestingly, and as mentioned, this AFS is the same as in Fig. 10. We showed a similar behavior in Fig. 12, when all the species were diluted at the original flat 50% dilution except for two species which were diluted independently at a range of values. The region was again split into one region where the system evolved towards the OFS, while in the other region, the system culminated in the same AFS as before. Once again, we saw that the system was most sensitive to the dilution of *m*_3_. Just as in Sec. I, we see that the model is much more sensitive to *m*_3_, presumably because it decreases by the greatest amount from its initial steady state to the OFS, which leads to greater sensitivity in its differential equation.

We also considered what would happen if we held all the species at a fixed 100% value while diluting only one of the species at a variety of levels and showed the results in Fig. 13. Interestingly, we found that if we diluted or enhance any single species in the direction of its OFS, that is where the system evolved while, if we diluted any species in the opposite direction of the OFS, it would instead evolve to the same AFS as before. We further noted that this study showed that the initial steady state was unstable to small perturbations and that we considered this to be an obvious shortcoming of this model. However, we discussed that artificially implementing a region of stability around the initial steady state would be arbitrary and that, instead, we should seek experimental dilution data within the stability region as well as outside in order to guide our models. In Fig. 14, we considered holding all but two species fixed at the initial steady state value and independently varying the dilution of the other two. We found a similar behavior to Fig. 13, where diluting in the direction of the OFS lead the system there and diluting in the other direction lead to the AFS. We also noted that the presence of an AFS and the near universality of it among these models, even though it was not part of the training data, nor was it explicitly programmed suggests that, with better data, we might predict alternate steady states which might be useful in the lab or the clinic and dilution regimens which might achieve them. In App. D, we presented various dilution scenarios for a single model for illustration.

As we have stated repeatedly throughout this note, the available data is not yet sufficient to make robust predictions and, therefore, the specific predictions made in this note should be taken as nothing more than illustrative of what sort of predictions we might make if we gathered better data. In particular, the data that would be the most helpful would be:

- Hourly measurements before dilution to determine the circadian cycle for the plasma constituents. As mentioned in [9], the behavior post-dilution could be impacted by the timing of the dilution events since they might be higher or lower in the morning relative to the evening. This would also give a more realistic model and understanding of the plasma system.
- Hourly measurements following dilution. Such data would have a large impact on disambiguating the various models that are only fit to the endpoints since it would constrain the models to fit the data in between the initial and final steady states.
- Measurements with a variety dilution amounts, including some infusions. This would further disambiguate the models and establish a region of stability around the steady states and lead to greater predictive power for new experiments.

We encourage the experimental community to carry out these experiments. We would also like to encourage the release of more precise concentration levels for each constituent, which would allow analysis beyond merely stating whether a concentration is relatively high or low. This could be important, in particular, in better understanding the catalyzed dynamics and their threshold values.

## Appendix A: The Structure of Our Dynamical System Equations

Modern systems [15] are built on the realization that the reactions that create and destroy the constituents of the system are intrinsically random and determined by probability. For the simplest example, a protein is typically unstable and will decay at some point. However, we do not know when it will decay. The best we can do is predict (or measure) the probability of decay per unit time. This is called the decay constant, let’s call it *λ*. With a large sample size, as we typically have in a biological sample, the approximate number of decays per unit time, then, is given by the expectation value for decays per unit time or, in other words, the probability per unit time multiplied by the number of species present, giving

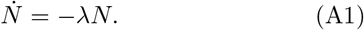

We see that the decay rate is linear.

On the other hand, some creation and/or destruction of a chemical species might be catalyzed and require the presence of one or more other species. For example, suppose the production of species 1 is catalyzed and requires the presence of species 2. Once again, given the probability per unit time that a single member of species 2 will produce species 1, the expected number of new copies of species 1 would be the product of the probability per unit time multiplied by the number of species 2. As a result, our first guess might be that the production of species 1 is linear in species 2. However, before we write a formula, we note that here the catalyzed production per unit time is typically not linear. For very small concentrations of species 2, it is approximately linear while there are plenty of free catalysts waiting for a member of species 2 to attach. However, as the density of species 2 increases, the catalyst eventually becomes saturated and the production of species 1 asymptotically approaches a constant value. This process is typically modeled with a decay probability per unit time of *g/*(*K*_2_ + *N*_2_), where *g* determines the maximal rate of production and *K*_2_ is roughly half the number of species 2 that saturates the catalyst. Multiplying by the concentration of species 2, we have a model of a catalyzed production of species 1 given by [16]

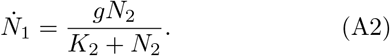

As constructed, this gives an approximately linear rate of (*g/K*_2_)*N*_2_ when *N*_2_ is small compared to *K*_2_ but an approximately constant maximal rate of *g* when *N*_2_ is large compared to *K*_2_ and a half maximal rate of *g/*2 when *N*_2_ = *K*_2_.

It is also possible that *N*_1_ catalyzes its own production, such that *N*_2_ is replaced by *N*_1_ in Eq. (A2). Additionally, *N*_1_ can catalyze its own destruction beyond its decay. In this case, we would have a negative sign, as in

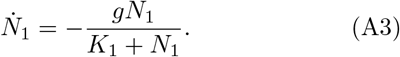

We cannot, on the other hand, have a catalyzed destruction of *N*_1_ that does not depend on the presence of *N*_1_, since *N*_1_ must be present to be destroyed. Therefore, we cannot have a negative sign in terms like Eq. (A2) where *N*_1_ does not appear on the right. In particular, it is this fact that removes the possibility of any species becoming negative. Any decrease of a species depends on a negative rate, but also depends on the presence of that species to occur. Therefore, with this requirement, the species in our models will not become negative.

Extending Eq. (A2), some creations that are catalyzed require the presence of more than one species. For example, a production catalyzed by two species can be modeled by

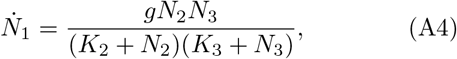

where *N*_2_ or *N*_3_ could also be *N*_1_. Moreover, a destruction catalyzed by two species could take the form

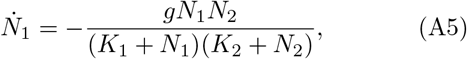

where at least one species on the right is *N*_1_. Extending this pattern to catalysts with greater numbers of species is possible, but increasingly less likely. We will only consider products of at most 2 species in this work.

Finally, the creation and destruction of species 1 could have contributions from many sources so, putting this all together, we have a system of equations like

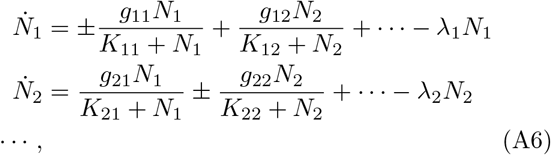

where the terms with products of species in a single term are in the ellipsis. As we can see, this set of equations can become quite complicated with a large set of unknowns. It would be quite difficult and nonlinear to determine all the constants, the *g*’s, *K*’s and *λ*’s.

In chemical systems studied elsewhere, the models have been analyzed in a narrow range of values for the constituents where each species has not traversed a significant range of its transition from low values (*N*_*i*_ ≪*K*_*i*_) to high values (*N*_*i*_ ≫*K*_*i*_). Therefore, in order to make calculations more efficient, *N*_*i*_*/*(*K*_*i*_ + *N*_*i*_) has been modeled as 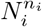, where *n*_*i*_ is some power typically between 0 and 2. Such a model is sometimes called an S-System model and written like [15]

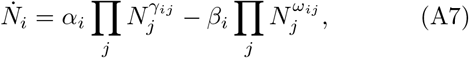

where *γ*_*ij*_ and *ω*_*ij*_ are the powers appropriate for the dynamics and fit to experimental data along with the rate constants *α*_*i*_ and *β*_*i*_. With this great simplification, fitting is more efficient but the meaning of the resulting equations is less clear.

Although this would be a simpler system, in the present context, where we are diluting the plasma to half or more of its original value, we expect the concentrations of the species to vary over a wide range including much of the transition region of each catalyst. We would also like to keep a description with greater meaning that might potentially give more insight into the dynamics. Therefore, we stick with the full Michaelis-Menten structure outlined in Eq. (A6).

The next thing we would like to do is split the species into four categories that we will call *m*_*i*_, *u*_*i*_, *h*_*i*_ and *c*_*i*_. Briefly, these categories have the following properties,

*m*_*i*_: These are the protein species that are measured in the plasma and undergo dilution.

*u*_*i*_: These are the protein species that are not measured, but are in the plasma and are diluted along with the measured species.

*h*_*i*_: These are species that are not measured and are not in the plasma and are thus not diluted. They are likely hidden inside the cells.

*c*_*i*_: These are species that are not measured and are not in the plasma and are also not diluted. Additionally, they are conserved and come in either the “excited” state or the “unexcited” state, which might represent the methylation status of a segment of deoxyribonucleic acid (DNA).

In the main body of this note, we will use *m*_*i*_, *u*_*i*_, *h*_*i*_ and *c*_*i*_ in place of the more generic *N*_*i*_. In the last category, the number of excited copies plus the number of unexcited copies adds to a total that does not change. Because of this, in principle, we would have two equations for each *c*_*i*_, one for the excited state and one for the unexcited state. For example, if we focus purely on *c*_1_ and call *c*_1_ and 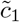 the excited and unexcited versions respectively, we might have

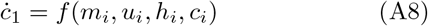

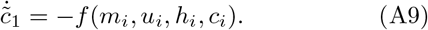

The symmetry between the two is required in order for

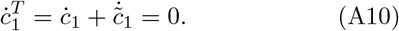

That is, this is required so that the total of the excited and unexcited states does not change. However, this symmetry enables us to remove one of the equations which is redundant. In particular, since 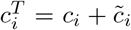 is constant, we can replace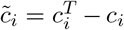. Since all our species concentrations will be taken as relative, we will define 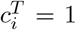and measure *c*_*i*_ as the relative value to the total. Therefore, we will use 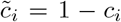 in this analysis in place of 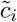. Moreover, since each catalyst can depend on either the excited version or the unexcited version, we will have terms in our differential equations of two types. Each term could either have

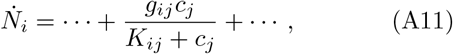

or,

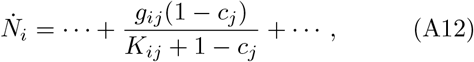

and similarly with product terms.

## Appendix B: Algorithmic Methods

Since neither the exact structure of the model nor the values of the parameters are known, we searched the space of models using a combination of simulated annealing and gradient descent to find the parameter values within a model structure and a genetic algorithm to find the optimal model structure. We will explain each below.

In the first step of the genetic algorithm, we created a random population of model structures. During the random creation of a model structure, we began by choosing the number of each type of species (the number of *m*_*i*_, *u*_*i*_, *h*_*i*_ and *c*_*i*_.) The number of *m*_*i*_ was always fixed by the number of observed species chosen from [1]. The minimum number of other species was an argument that was specified by the user. In this stage, the algorithm chose a number of the other species exactly equal to the minimum number requested, but split them randomly between the *u*_*i*_, *h*_*i*_ and *c*_*i*_.

Once the number of each species was determined for a model, the algorithm cycled through the equation for each species and randomly chose a number of terms between 2 and 6 for the right side of the equation. It then cycled through each of these terms. For each term, it randomly chose whether it only depended on one species or two. It then randomly chose which species was in each term, choosing either one species or two for each term. If either species was conserved, it randomly chose whether it used the excited state (and used *c*_*i*_) or the unexcited version (and used 1− *c*_*i*_). This completed the general structure. The final piece to make the model functional was to randomly generate the parameters with values between 0 and 1. If the term was allowed to have a negative coefficient [see Eqs. (A3) and (A5)], we randomly chose whether to flip the sign of the coefficient.

Given a new model structure, the next step is to regress the parameters to optimize the fit to the data. The data is a series of measurements with an intermediate dilution. In the present case, it is a set of two time points. The measured species were observed before the dilution, the dilution occurred and the species were measured at a later time after a steady state was reestablished. Although the before and after species concentrations may not strictly be constant (they likely vary cyclically each day and depend on daily nutrition and other factors), in the absence of sufficient data, we took them to be constant for the purpose of illustrating what could be achieved with these models.

Given a model with a set of parameters, we began by solving for a set of initial values for the species that was constant (where the right side of the equations was zero.) We did this by using the scipy.optimize Python package by scanning over initial values in a grid and finding nearby roots. With these roots in hand, which are constant steady states, we diluted each one by the dilution amount in the data. We then used the Stimator Python package [17] to solve the differential equation and find the time series after the dilution.

For each initial steady state tested, there were several checks performed on the resulting time series. If the results failed any of the checks, the initial steady state was discarded and the next one was tested. The checks are as follows: the final species concentrations were tested to be a steady state, both pairs of steady states were tested to not be trivial (the measured values could not be equal to zero), the concentrations were tested to never become negative, any conserved species were tested never to exceed 1, the final steady state was tested to not be identical to the initial steady state (we didn’t want the dilution to return the system to the original steady state).

For each model and parameter point, each initial steady state that passed the previous tests were scored for their proximity to the experimental data. For each measured species, the following was calculated:

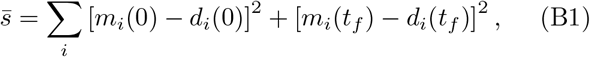

where *t*_*f*_ is the final time, *d*_*i*_ is the value from data for this species and *m*_*i*_(0) and *d*_*i*_(0) were before the dilution. We then multiplied this by one plus the number of terms in the equations and multiplied by one plus the number of extra species used. The purpose of this was to favor models that were more economical in the use of terms and extra unmeasured species. Finally, we took the square root to obtain the final score,

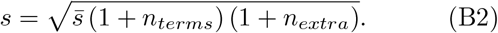

This score served the basis of comparing models. A model with a lower score was a better fit and more optimized to the data.

Since our parameters are non-linear and of high dimension, we chose the method of simulated annealing to efficiently search the space. Given a parameter, perhaps *g*_31_, and a current best value for that parameter, we choose a new value with a flat random distribution within a radius. The radius begins at a large value and diminishes slowly to a vanishingly small value. We found that a simple linear reduction in this radius worked well. In the beginning, when we are not confident we are near an optimal parameter point we search a wide region of the parameter space. However, as our search progresses and our parameters approach closer to a locally optimal solution, we search a smaller and smaller region around that parameter point looking for an even better solution.

We further augmented our search by using a gradient descent method for each parameter point randomly chosen by the simulated annealing function. In particular, for each random parameter point, we calculated the gradient in the parameter space around that point and then followed the gradient down towards more optimal solutions. We found that coupling these two methods performed better than either method in isolation. We note that these two methods do not necessarily produce a globally optimized solution. They only find an optimized solution within a local region of parameter space and there still may be other more optimal parameter points within the same model structure.

Once a model is locally optimized, we add it to the genetic population if its model score is lower than the worst score in the population. If we add it, we remove the worst score to keep the population size the same. Once we have filled the requested population size, we begin producing new models in a few different ways. We randomly choose among the methods of producing new models. Twenty percent of the time, we simply create a brand-new random model as above. Forty percent of the time, we choose two models from the population with the same number of each type of species (the same number of measured *m*_*i*_, unmeasured *u*_*i*_, hidden *h*_*i*_ and conserved *c*_*i*_ species) and mate them. This consists in producing a new model with the same species but with a mixture of terms coming from the two parents. Ten percent of the time, we choose a model from the population and add a new species from among the *u*_*i*_, *h*_*i*_ and *c*_*i*_ and a random set of terms for its DEs. Ten percent of the time, a model is chosen from the population and a new term is added to one of its DEs. Ten percent of the time, a model from the population has a species from among *u*_*i*_, *h*_*i*_ and *c*_*i*_ removed. Ten percent of the time, a model from the population has a term removed from one of the DEs. If any of these fail, a new attempt to create a new model is tried again.

Finally, a central program starts jobs on each cpu of a parallel machine and monitors the progress of each job, collecting its results, inserting it in the population if appropriate and starting new jobs. Each job runs in parallel on its own thread, using the Condor cluster software [18].

## Appendix C: Minimal Models

In this appendix, we will present plots and differential equations for a selection of representative models. Although we collected a population of a thousand such models, we found that they could be organized into the following three groups depending on their behavior following dilution. They are

Group 1: *m*_0_, *m*_1_ and *m*_2_ began rising while *m*_3_ began decreasing immediately following dilution.

Group 2: All of *m*_0_, *m*_1_, *m*_2_ and *m*_3_ began increasing immediately following dilution.

Group 3: Same as Group 1, but with several oscillations before coming to a steady state.

Of our population of a thousand models, 460 were of class 1, 528 were of class 2 and 12 were of class 3. Within each class, the models differed in how quickly they approached the final steady state as well as how far they overshot before returning to the steady state.

### 1. Class 1 Models

There were a small number of models from this group where *m*_0_, *m*_1_ and *m*_3_ evolved monotonically towards the final steady state while only *m*_2_ had one or two peaks before settling down towards its steady state. We show an example of such a model in Fig. 15(a). It has differential equations,

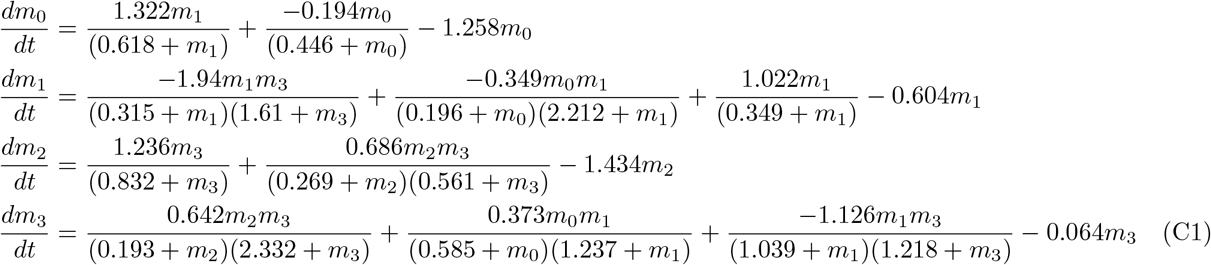

The other models in this group were not monotonic in any of the species. They all overshot before approaching the final steady state. *m*_0_, *m*_1_ and *m*_3_ usually only had a significant overshoot once while *m*_2_ typically overshot twice, before attaining the steady state. However, within that group, the height and depths of the peaks as well as the timing differed. We chose three models to illustrate this.

The plot of the first of these models can be found in Fig. 15(b) and has differential equations,

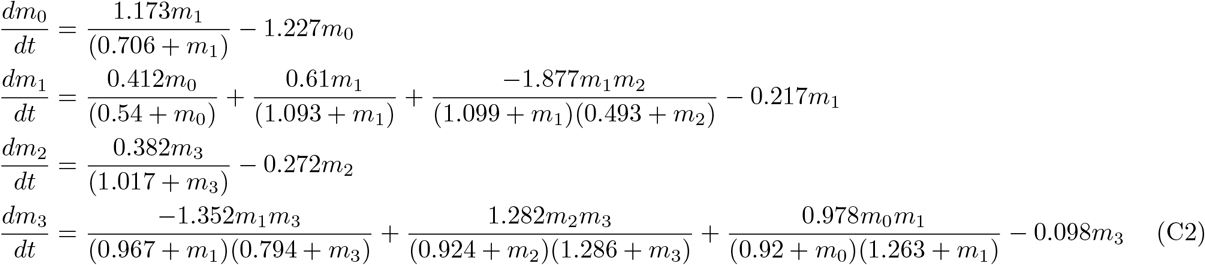

The plot of the second of these models can be found in Fig. 15(c) and has differential equations,

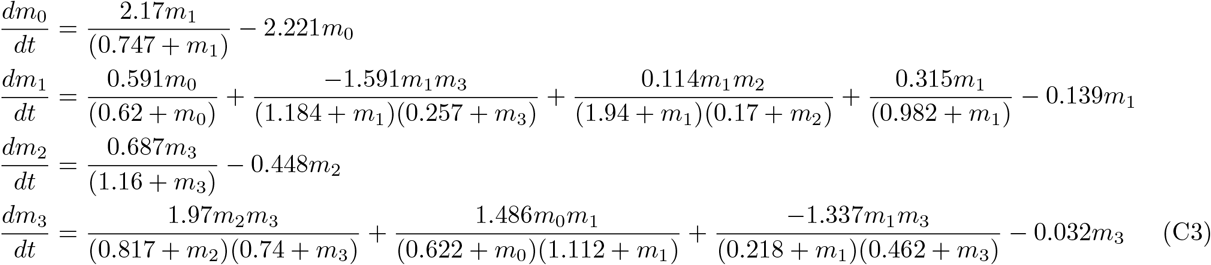

The plot of the thrid of these models can be found in Fig. 15(d) and has differential equations,

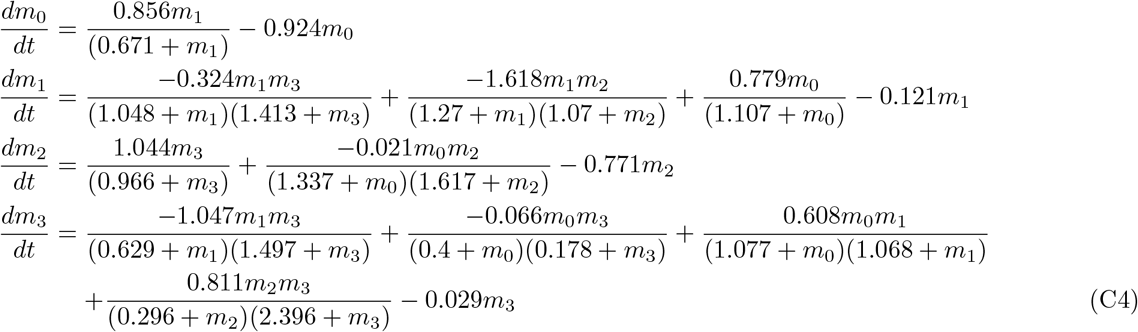

### 2. Class 2 Models

All the models in this group are qualitatively similar. The species *m*_0_ and *m*_1_ all monotonically increase to their final steady state, while *m*_2_ and *m*_3_ increase to a peak and then fall monotonically to their respective final steady states. In Fig. 16, we show the plots for 4 of these models and describe their differential equations here.

The plot for the first illustrative model of this class can be found in Fig. 16(a) and has differential equations,

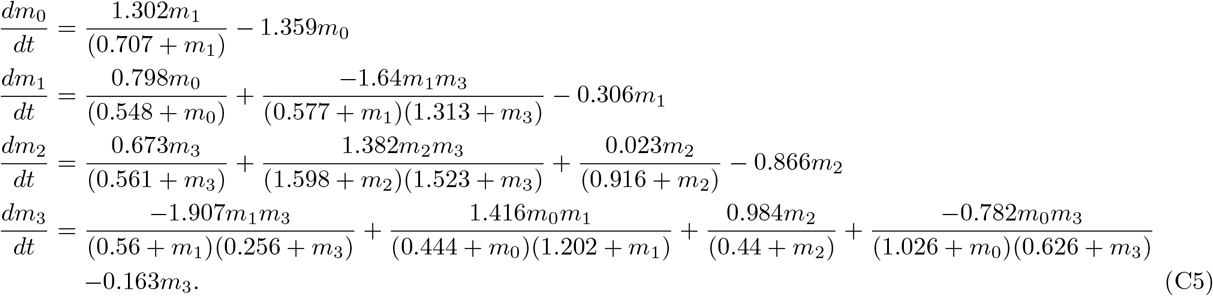

The plot for the second illustrative model of this class can be found in Fig. 16(b) and has differential equations,

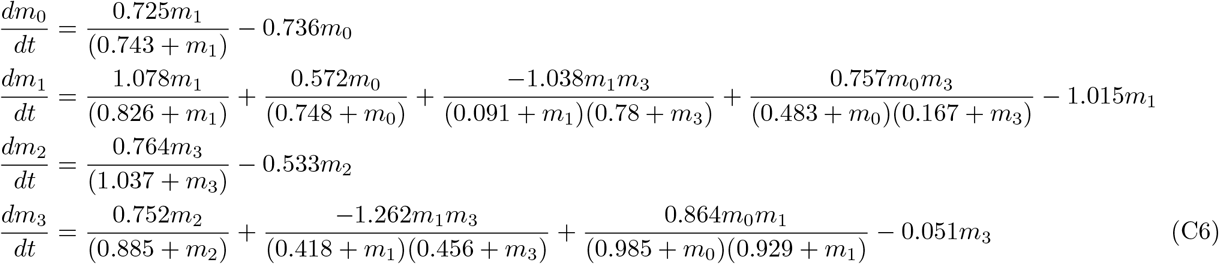

The plot for the third illustrative model of this class can be found in Fig. 16(c) and has differential equations,

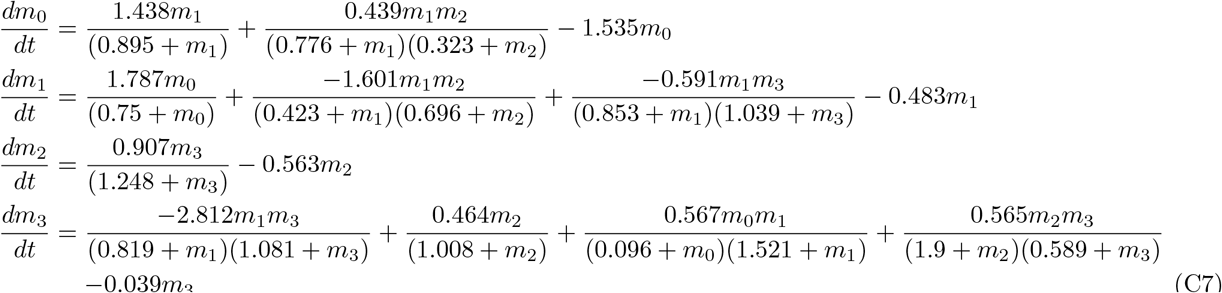

The plot for the fourth illustrative model of this class can be found in Fig. 16(d) and has differential equations,

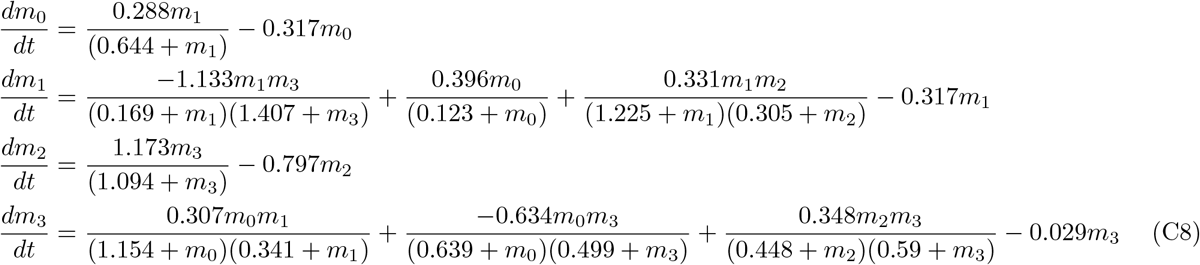

### 3. Class 3 Models

Our final class of models has an oscillatory pattern after dilution. In this group, *m*_0_ and *m*_1_ shoot up after dilution, but then they oscillate around the final steady state until the oscillatory energy is dissipated and they relax to their final steady state. *m*_2_ first rises, before turning down and entering an oscillatory pattern until it relaxes to its final steady state. *m*_3_ falls into oscillation, finally approaching its final steady state. Plots for four illustrative models can be seen in Fig. 17.

The plot of the first example model from this group can be found in Fig. 17(a) and has differential equations,

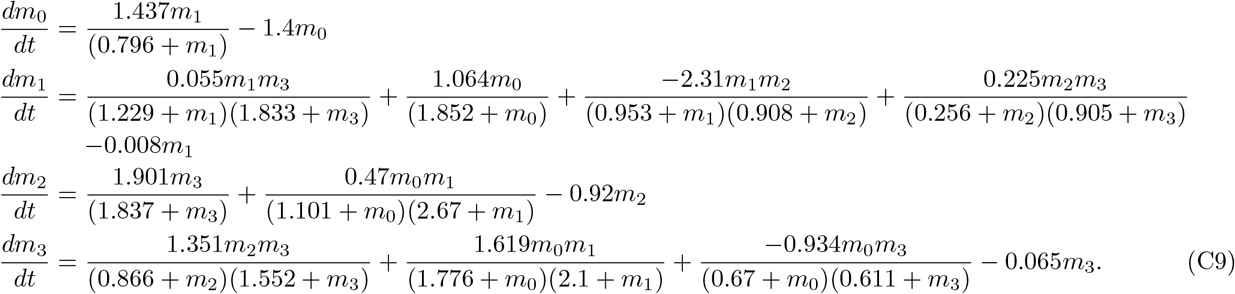

The plot of the second example model from this group can be found in Fig. 17(b) and has differential equations,

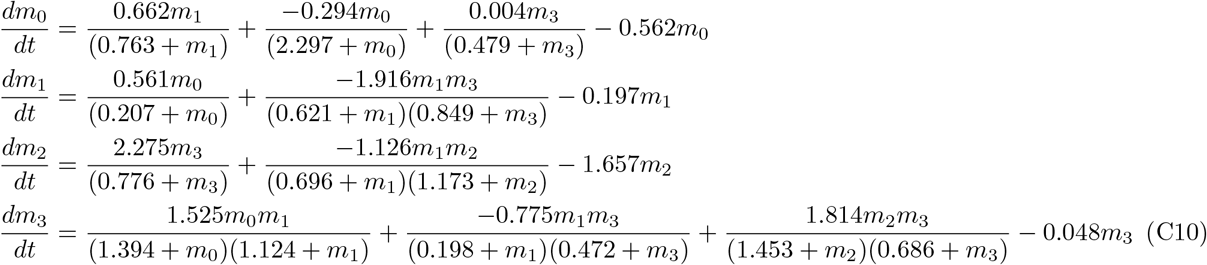

The plot of the third example model from this group can be found in Fig. 17(c) and has differential equations,

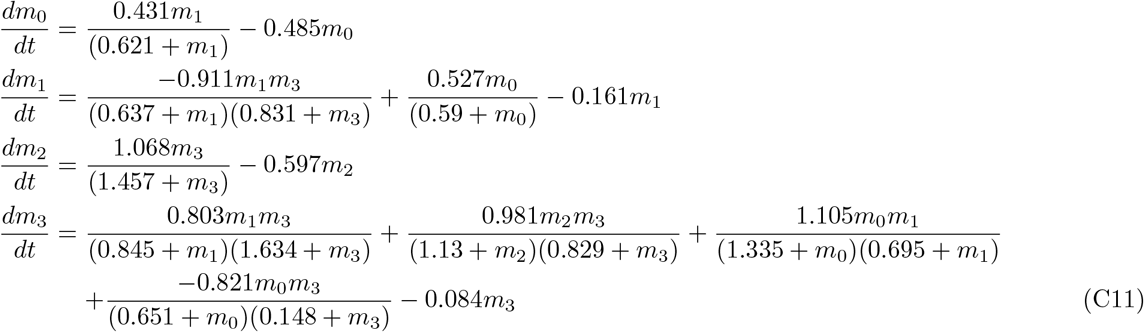

The plot of the fourth example model from this group can be found in Fig. 17(d) and has differential equations,

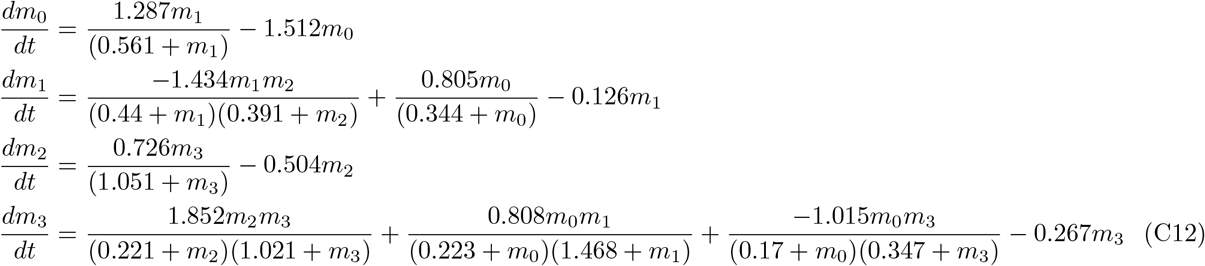

## Appendix D: Modified Dilution Examples

In this appendix, we present a few examples of the transition from the original final steady state to a new final state when we change the dilution of one or more constituents.

In Fig. 18, we present the response of the model in Fig. 17(d) and Eq. C12 to three scenarios. In the first column, we consider a flat 50% dilution of *m*_0_, *m*_1_ and *m*_2_, but a varying dilution or enhancement of *m*_3_. We can see that for dilutions below approximately 1.0 (a non-dilution), the constituents still evolve to the original final steady state (OFS – top plot), as shown on the right of Eq. (6). However, once enhancements begin, in this example, the species instead evolve to an alternate steady state (ASS – bottom plot) with final values 0.17, 0.14, 1.02 and 2.50. In the second column, *m*_0_, *m*_1_ and *m*_2_ are held at the pre-dilution value but *m*_3_ is allowed to have a variety of dilution values. Similarly to the left column, diluting *m*_3_ moves it in the OFS direction and that is where it evolves towards (top plot) and, when it is enhanced, it evolves to the same AFS as before (bottom plot). In the third column, we instead hold *m*_1_, *m*_2_ and *m*_3_ while varying the dilution of *m*_0_. Unlike the previous two cases, a dilution of *m*_0_ is a move in the direction opposite the OFS and so it evolves towards the AFS (top plot) while an enhancement is towards the AFS and that is where it ends up (bottom plot). We also see in the middle plot of the right two columns that not diluting *m*_3_ or *m*_0_, respectively, causes the state to remain in the original state and not change. This is expected since none of the constituents are changed.

**FIG. 18.**
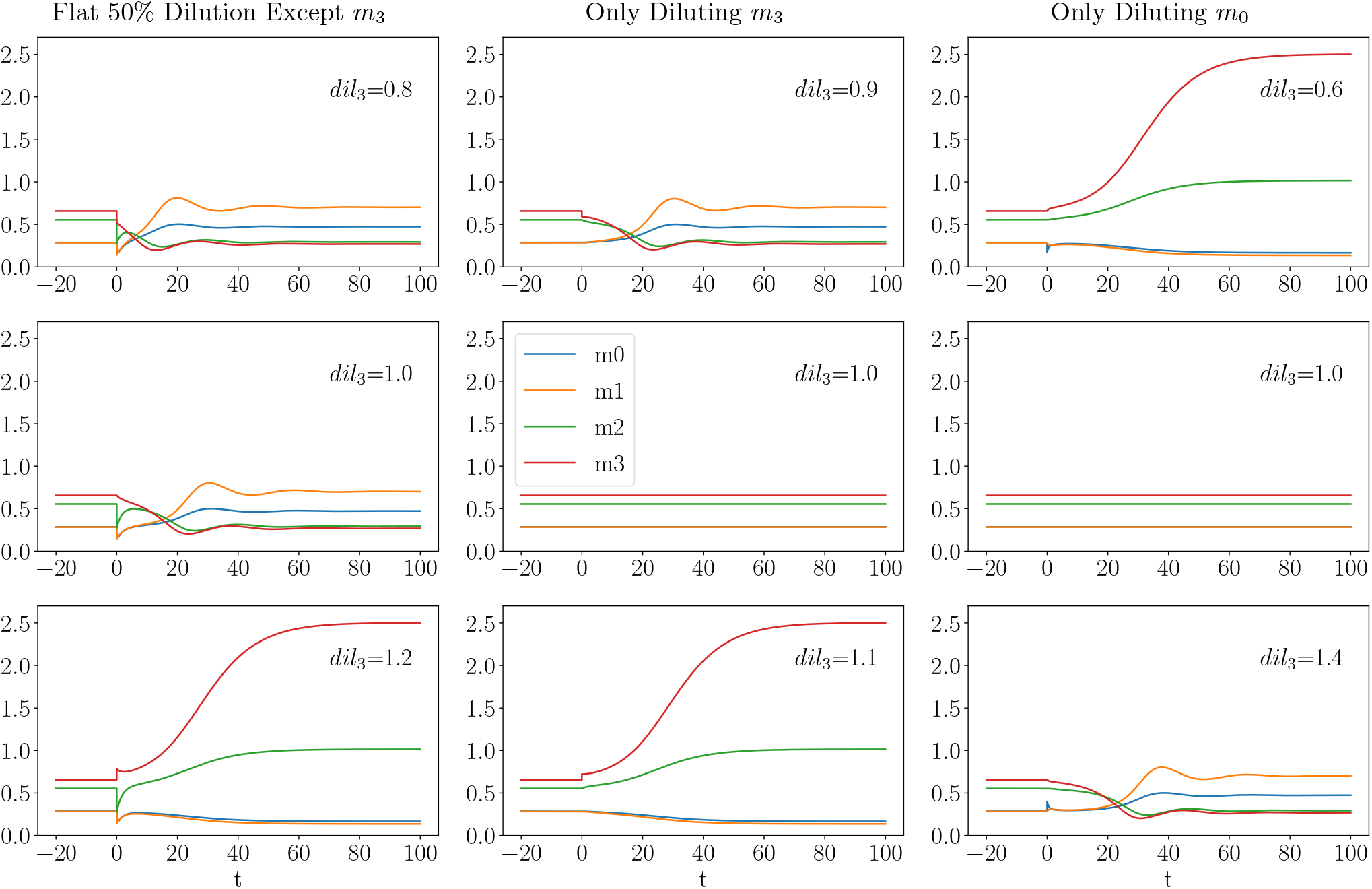
The response of the model presented Fig. 17(d) under three scenarios. The first column presents what happens when *m*_0_, *m*_1_ and *m*_2_ are all diluted at 50%, but *m*_3_ is diluted or enhanced by a different value, the second and third columns present the result of holding *m*_0_, *m*_1_ and *m*_2_ at their predilution value and diluting or enhancing *m*_3_ by a different value. The dilution of *m*_3_ is given in the top right with *dil*_3_.

Behavior similar to this, although with different transition dilution values, different evolution details and different alternate final steady states, is expected in a large number of models based on the analysis in Sec. II, especially that surrounding Fig. 11, especially the bottom panel, and Fig. 13. In fact, that is what we find, generally speaking. We chose this particular model because the AFS was not very high so we could show the behavior over many different dilutions in a single figure and it had interesting evolution properties. As noted in Sec. II and here, the models are most sensitive to *m*_3_ in the case that the other constituents are held at a flat 50% dilution and that is interesting. It suggests that a dilution followed by an enhancement of one of the plasma constituents that previously fell substantially might be enough to move the system into a different final state. Moreover, the system is very sensitive to even a single species changing while the others are not diluted at all. In fact, we see, at least in these very limited models, that the enhancement of a single species is sufficient to drive the system into either the OFS or AFS, depending on which species is chosen. On the other hand, we consider it important to remember that these models are still extremely limited by the lack of sufficient data. Given greater data which force the models into more complex structures with hidden constituents could change this behavior. Nevertheless, the fact that his behavior emerges in a large number of the models even though it was not built in suggests that this general behavior should be considered in future studies of dynamical models for the plasma system.

## References

[1] Mehdipour, M., Skinner, C., Wong, N., Lieb, M., Liu, C., Etienne, J., Kato, C., Kiprov, D., Conboy, M. J., & Conboy, I. M. (2020). Rejuvenation of three germ layers tissues by exchanging old blood plasma with saline-albumin. Aging, 12(10), 8790–8819. https://doi.org/10.18632/aging.103418

[2] Mehdipour, M., Mehdipour, T., Skinner, C.M. et al. Plasma dilution improves cognition and attenuates neuroinflammation in old mice. GeroScience 43, 1–18 (2021). https://doi.org/10.1007/s11357-020-00297-8

[3] Conboy IM, Conboy MJ, Wagers AJ, Girma ER, Weissman IL, Rando TA. Rejuvenation of aged progenitor cells by exposure to a young systemic environment. Nature. 2005; 433:760–64. https://doi.org/10.1038/nature03260 PMID:15716955

[4] Rebo J, Mehdipour M, Gathwala R, Causey K, Liu Y, Conboy M, & Conboy I. A single heterochronic blood exchange reveals rapid inhibition of multiple tissues by old blood. Nature Communications 1, 13363 (2016). https://doi.org/10.1038/ncomms13363

[5] Kiprov DD. Therapeutic apheresis delivery systems in the US. Transfus Apher Sci. 2003; 28:163–64. https://doi.org/10.1016/s1473-0502(03)00005-3 PMID:12679121

[6] Kiprov DD, Golden P, Rohe R, Smith S, Hofmann J, Hunnicutt J. Adverse reactions associated with mobile therapeutic apheresis: analysis of 17,940 procedures. J Clin Apher. 2001; 16:130–33. https://doi.org/10.1002/jca.1024 PMID:11746539

[7] Kiprov DD, Hofmann JC. Plasmapheresis in immunologically mediated polyneuropathies. Ther Apher Dial. 2003; 7:189–96. https://doi.org/10.1046/j.1526-0968.2003.00028.x PMID:12918942

[8] Mehdipour M, Etienne J, Liu C, Mehdipour T, Kato C, Conboy M, Conboy I, Kiprov DD. Attenuation of age-elevated blood factors by repositioning plasmapheresis: A novel perspective and approach. Transfus Apher Sci. 2021 Jun;60(3):103162. doi: 10.1016/j.transci.2021.103162. Epub 2021 May 21. PMID: 34083162.

[9] Flickenger Rob. Plasma donation induces a protein expression profile shift in circulating human blood. bioRxiv. doi:10.1101/2021.10.30.466597.

[10] Katsimpardi Lida et al. Vascular and Neurogenic Rejuvenation of the Aging Mouse Brain by Young Systemic Factors. Science 344, 6184 (2014) pgs. 630–634. DOI: 10.1126/science.1251141

[11] Dmytro Shytikov, Olexiy Balva, Edouard Debonneuil, Pavel Glukhovskiy, and Iryna Pishel. Aged Mice Repeatedly Injected with Plasma from Young Mice: A Survival Study. BioResearch Open Access 3, No. 5 (2014). DOI: 10.1089/biores.2014.0043

[12] Horvath, Steve et al. Reversing age: dual species measurement of epigenetic age with a single clock. bioRxiv. doi: 10.1101/2020.05.07.082917

[13] Suh, M.R. et al. Efficacy of cord blood cell therapy for Hutchinson-Gilford Progeria Syndrome – A case report. International Journal of Molecular Sciences (2021), 22(22), 12316. DOI: 10.3390/ijms222212316

[14] Boada, M. et al. Neuropsychological, neuropsychiatric, and quality-of-life assessments in Alzheimer’s disease patients treated with plasma exchange with albumin replacement from the randomized AMBAR study. Alzheimer’s Dementia (2021). DOI: 10.1002/alz.12477

[15] For a nice review, see Voit E.O. and Ferreira A.E.N. Computational Analysis of Biochemical Systems: A Practical Guide for Biochemists and Molecular Biologists. Cambridge University Press. 2000.

[16] Michaelis L, Menten MM. The kinetics of invertin action. 1913. FEBS Lett. 2013 Sep 2;587(17):2712–20. doi: 10.1016/j.febslet.2013.07.015. Epub 2013 Jul 15. PMID: 23867202.

[17] Ferreira António. S-timator: dynamical systems modelling in python [Internet]. c2006–2015 [cited 2021 Nov 22]. University of Lisbon. Available from: https://webpages.ciencias.ulisboa.pt/aeferreira/stimator/index.html

[18] Thain Douglas, Tannenbaum Todd and Livny Miron. Distributed computing in practice: the Condor experience. Concurrency - Practice and Experience, 17, 2–4 (2005), pgs 323–356.

